# Nuclear envelope-vacuole contacts mitigate nuclear pore complex assembly stress

**DOI:** 10.1101/2020.03.23.001719

**Authors:** Christopher L. Lord, Susan R. Wente

## Abstract

The intricacy of nuclear pore complex (NPC) biogenesis imposes risks of failure that can cause defects in nuclear transport and nuclear envelope morphology, however, cellular mechanisms utilized to alleviate NPC assembly stress are not well-defined. In the budding yeast *Saccharomyces cerevisiae*, we demonstrate that *NVJ1*- and *MDM1*-enriched nuclear envelope (NE)-vacuole contacts increase when NPC assembly is compromised in several *nup* mutants, including *nup116ΔGLFG* cells. These interorganelle nucleus-vacuole junctions (NVJs) cooperate with lipid droplets to maintain viability and enhance NPC formation in assembly mutants. Additionally, NVJs function with *ATG1* to promote vacuole-dependent remodeling in *nup116ΔGLFG* cells, which also correlates with proper NPC formation. Importantly, NVJs significantly improve the physiology of NPC assembly mutants, despite having only negligible effects when NPC biogenesis is unperturbed. Collectively, these results define how NE-vacuole interorganelle contacts coordinate responses to mitigate deleterious cellular effects caused by disrupted NPC assembly.

**Summary:** How cells respond to deleterious effects imposed by disrupted nuclear pore complex (NPC) assembly are not well-defined. The authors demonstrate nuclear envelope-vacuole interactions expand in response to perturbed NPC assembly to promote viability, nuclear envelope remodeling, and proper NPC biogenesis.

## Introduction

The nuclear envelope (NE) forms a semi-permeable barrier surrounding genomic DNA, with its two lipid bilayers fused at points where nuclear pore complexes (NPCs) are embedded. As multi-subunit assemblies composed of nucleoporins (Nups), NPCs are essential for nucleocytoplasmic transport. A set of scaffolding Nups provide a core structural platform on which FG and GLFG Nups assemble and serve to facilitate the transport of different types of cargos (Beck and Hurt, 2017). FG Nups contain domains rich in intrinsically disordered phenylalanine-glycine (FG) repeats (Denning et al., 2003; Hough et al., 2015), whereas GLFG domains are a subset that contain glycine-leucine-phenylalanine-glycine repeats (Wente et al., 1992). Both FG and GLFG Nups transiently interact with specific nuclear transport receptors (NTRs) to allow movement of NTR-cargo complexes through NPCs (Strawn et al., 2004).

NPC assembly is an elaborate process, particularly when NPCs are inserted into the intact NE during interphase. Unlike metazoan cells, which also re-form NPCs when the NE is reconstituted after cell division, *S. cerevisiae* cells undergo a closed mitosis and therefore only directly assemble NPCs into the NE. Disrupting the function of several Nups (Wente and Blobel, 1993; Chadrin et al., 2010; Makio et al., 2009; Boehmer et al., 2003; Aitchison et al., 1995; Doye et al., 1994; Onischenko et al., 2017), the GTPase Ran (Ryan et al., 2003; Walther et al., 2003), and NTRs (Ryan et al., 2007; Lusk et al., 2002) inhibits NPC formation, suggesting a functional nuclear transport system is required for further NPC production. Reticulons (Dawson et al., 2009), lipid regulating proteins (Hodge et al., 2010; Zhang et al., 2018), and Torsins (Laudermilch et al., 2016; Pappas et al., 2018; VanGompel et al., 2015) are also involved, likely by regulating properties of the NE to promote NPC biogenesis. Additionally, Vps4 and Heh1/2 cooperatively function as a surveillance system that promotes the clearance of misassembled NPCs in a Chm7-dependent manner (Webster et al., 2014, 2016; Thaller et al., 2019).

The NE in *S. cerevisiae* cells contacts vacuoles at regions termed nucleus-vacuole junctions (NVJs) (Pan et al., 2000; Kvam and Goldfarb, 2006). Although morphologically distinct, vacuoles are functionally similar to metazoan lysosomes, serving as acidic organelles involved in protein degradation and nutrient storage. Several outer NE membrane proteins including Nvj1 (Pan et al., 2000) and Mdm1 (Henne et al., 2015) are present at NVJs and tether the NE to the vacuole through specific interactions with corresponding vacuolar proteins like Vac8 (Pan et al., 2000; Jeong et al., 2017) or phospholipid species such as phosphatidylinositol 3-phosphate (Henne et al., 2015). NVJs expand when cells are grown using carbon sources other than glucose (Hariri et al., 2018; Bean et al., 2018) or to promote piecemeal autophagy of the nucleus (PMN) in response to nutrient starvation (Roberts et al., 2002), which involves autophagy-dependent degradation of specific nuclear proteins in the vacuole (Krick et al., 2008). Proteins that promote lipid droplet (LD) formation are also enriched at NVJs (Kohlwein et al., 2001), which serve as non-exclusive sites for LD synthesis. LDs are required for the viability of NPC assembly-defective *brr6-1* mutant cells (Hodge et al., 2010), suggesting LDs and NPCs are functionally linked.

While an ample amount of research has defined a set of models for how NPCs form, there is a significant lack of information about how cells respond to sustained interphase NPC assembly defects; such defects often manifest as clusters of misassembled intermediates that herniate the NE (Thaller and Lusk, 2018; Laudermilch et al., 2016) and disrupt nuclear transport and growth (Wente and Blobel, 1993; Aitchison et al., 1995; Hodge et al., 2010). Assembly defects are not only caused by inherent complexities associated with NPC biogenesis, but are also observed in replicatively aged *S. cerevisiae* cells (Rempel et al., 2019) as well genetic models of early-onset torsion dystonia (Laudermilch et al., 2016; Pappas et al., 2018). Here we investigate *S. cerevisiae* mutants with altered NPC assembly as models to determine how cells respond to NPC assembly stress. We find that NPC assembly mutants exhibit dramatic expansion of *NVJ1*- and *MDM1*-dependent NE-vacuole contacts and increased levels of LDs. Furthermore, NVJs and LDs function in a cooperative manner to promote viability and NPC formation in NPC assembly mutants. NE-vacuole contacts also function with autophagy factors to remodel the NE when NPC assembly is disrupted, which also results in fewer misassembled NPCs. Collectively, we reveal novel mechanisms coordinated by NE-vacuole interactions that promote proper NPC formation when NPC biogenesis is at least partially compromised.

## Results

### Nup116’s GLFG domain is required for NPC assembly and modulates NE-vacuole interactions

Mutant *nup116ΔGLFG* cells in a W303 strain background display NPC biogenesis defects in the absence of Nup188 (Onischenko et al., 2017), indicating that Nup116’s GLFG domain is required for NPC assembly when other factors are compromised. To further ascertain how Nup116’s GLFG domain regulates NPC formation, a *GFP-nic96 nup116ΔGLFG* strain produced in an S288C background was analyzed for GFP-Nic96 foci (indicative of NPC assembly defects) at temperatures ranging from 25 to 36°C. *GFP-nic96 nup116ΔGLFG* mutants displayed temperature-dependent increases in NPC foci formation, with ∼50% of these cells containing foci at 36°C compared to ∼10% of *GFP-nic96* cells (Fig. 1, A and B). Although NPC clustering in *GFP-nic96 nup116ΔGLFG* cells was more pronounced at 36°C, significant levels of foci were present at 30 and 34°C. Inhibiting new protein synthesis with cycloheximide prevented foci formation when mutants were shifted from 25 to 36°C (Fig. S1 A), indicating that the foci resulted from disrupted *de novo* NPC assembly.

**Figure 1:**
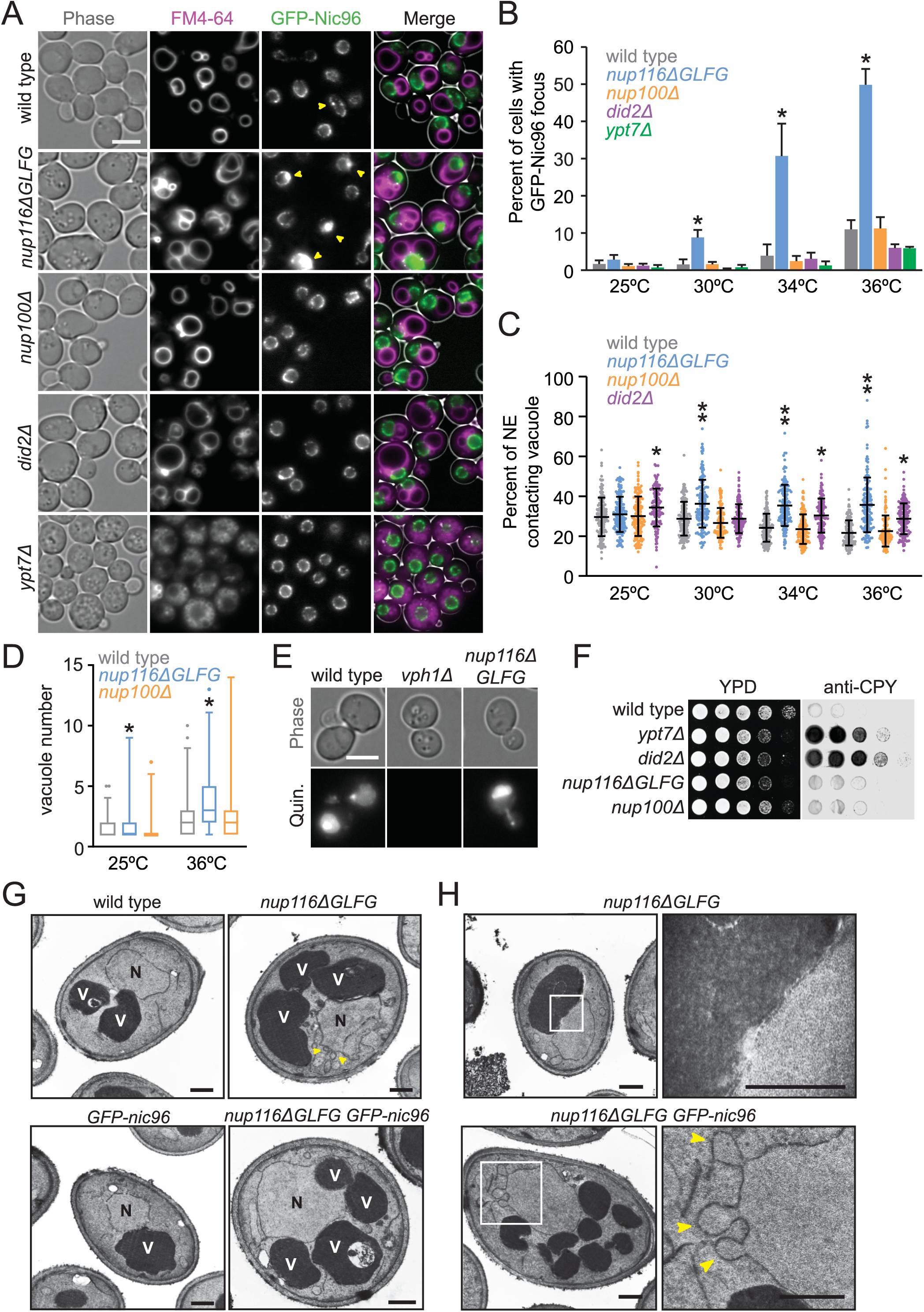
*nup116ΔGLFG* mutants display increased levels of GFP-Nic96 clustering and NE-vacuole contacts. **(A)** FM4-64-stained *GFP-nic96* yeast grown at 36°C. Arrows point to clustered NPC foci. “Phase” refers to phase contrast imaging. Bar, 5 μm. **(B)** Percentage of cells with at least one GFP-Nic96 focus at temperatures ranging from 25 – 36°C. Asterisk indicates a p-value of less than 0.05 when compared to wild type at the same temperature using an unpaired two-tailed Student’s t-test with an n of at least 100 cells from 3 independent experiments. **(C)** Percentage of NE contacting FM4-64-labeled vacuole membranes (detailed in methods); each dot represents value from one cell. Forty cells were analyzed from 3 independent experiments at temperatures ranging from 25 – 36°C. One asterisk indicates a p-value of less than 0.001 when compared to wild type at the same temperature using Dunn’s post-hoc test; two asterisks indicate a p-value of less than 0.03 when also compared to *did2Δ* contacts. **(D)** Vacuole number per cell; asterisk indicates a p-value of less than 0.01 when compared to wild type at the same temperature using a two-tailed Mann-Whitney U test calculated from over 30 cells from 3 independent experiments. Box and whiskers represent 1-99 percentile. **(E)** Quinacrine-stained cells to assess vacuole acidity. Quinacrine fluorescence indicates vacuoles are acidic; *vph1Δ* cells served as a positive control for vacuoles lacking acidity (Preston et al., 1989). **(F)** Yeast were serially diluted onto 2 YPD plates and grown at 28°C; one was covered with nitrocellulose to absorb secreted proteins and blotted with anti-CPY. Secreted CPY results from blocks in Golgi-vacuole transport (Robinson et al., 1988) **(G)** TEM images of listed strains at 36°C. Nuclei are labeled with a black “N”, while vacuoles are labeled with a white “V”. Arrows point to NE herniations. Bars, 500 nm. **(H)** TEM images of *nup116ΔGLFG* cells at 36°C zoomed into regions of NE-vacuole contacts (top) or NE herniations (bottom). Arrows point to NE herniations. Bars, 500 nm.

Kap121 is one of several NTRs whose transport is regulated by Nup116’s GLFG domain (Strawn et al., 2004; Lord et al., 2015; Terry and Wente, 2007). If foci were forming in the *GFP-nic96 nup116ΔGLFG* cells due to disrupted Kap121-dependent transport, we predicted that *GFP-nic96 kap121* mutants would also show foci formation. Consistent with this hypothesis, *kap121-34* and *kap121-41* mutants displayed increased GFP-Nic96 foci formation at the semi-permissible temperature of 32°C, while *kap122Δ* and *kap114Δ* mutants had no effect (Fig. S1, B and C). We therefore concluded that deletion of Nup116’s GLFG domain in S288C cells causes temperature-dependent NPC assembly defects that may result from compromised Kap121-dependent nucleocytoplasmic transport.

Given their relatively modest growth defects (Fig. S1 D), *nup116ΔGLFG* mutants were leveraged to determine how perturbed NPC assembly affects other aspects of cell function. Phase contrast images of *GFP-nic96 nup116ΔGLFG* cells indicated possible changes in vacuole morphology; the endomembrane-staining dye FM4-64 confirmed these mutants exhibited elevated contacts between the NE and vacuole membrane (Fig. 1, A and C; Fig. S2) and significantly more vacuoles per cell (Fig. 1 D). Increased NE-vacuole interactions were particularly evident at higher temperatures that correlated with disrupted NPC assembly. Mutant cells lacking the GLFG Nup100 (*nup100Δ*) displayed neither increased GFP-Nic96 foci nor altered vacuole morphology/number, indicating these types of defects are not caused by all *nup* mutants. The morphological changes in *nup116ΔGLFG* vacuoles did not appear to significantly impact vacuolar function, as they remained acidic (Preston et al., 1989) and the peptidase CPY was properly transported to the vacuole from the Golgi (Robinson et al., 1988) (Fig. 1, E and F).

Importantly, *did2Δ* and *ypt7Δ* cells, which cause vacuole fragmentation and transport defects (Ohsumi et al., 1988), showed no increase in GFP-Nic96 foci formation, indicating vacuole number does not impact NPC assembly (Fig 1, A and B). Due to severe vacuole fragmentation in *ypt7Δ* cells, NE-vacuole interactions were not quantified; *did2Δ* mutants displayed increased NE-vacuole contacts at most temperatures tested, but unlike *nup116ΔGLFG* cells, the phenotype was not exacerbated in a temperature-dependent manner (Fig. 1, A and C). TEM images of *nup116ΔGLFG* mutants grown at 36°C confirmed extensive direct contacts between the NE and vacuole membranes (Fig. 1, E and F). NE herniations were frequently detected in *nup116ΔGLFG* cells at 36°C, and were structurally similar to those observed in *nup116Δ* mutants (Wente and Blobel, 1993). Overall, at elevated temperatures, *nup116ΔGLFG* cells exhibited NPC biogenesis defects that correlated with increased physical interactions between the NE and vacuole membrane that did not impact vacuole function.

### NE-vacuole interactions expand in other NPC assembly mutants

Based on the lack of NPC foci and vacuole morphology phenotypes in *nup100Δ* cells, we hypothesized that increased NE-vacuole contacts could result from disrupted NPC assembly or associated NE morphological changes (i.e., herniations). NE-vacuole interactions were therefore measured in *nup120Δ* and *nup133Δ* mutants at 30°C, a temperature at which both strains are viable, yet also exhibit severe NPC assembly defects (Aitchison et al., 1995; Doye et al., 1994). The predicted integral NE/ER protein Pho88-GFP (Aviram et al., 2016) was used a marker because the NPC clustering defects are so strong in *nup120Δ* and *nup133Δ* mutants that most of their nuclear rim areas are devoid of NPCs (Aitchison et al., 1995; Doye et al., 1994). FM4-64 labeling revealed extensive contacts between the NE and vacuoles in both mutants (Fig. 2, A and B). Together, this data supported a model whereby mutations that perturb NPC assembly and NE integrity also promote changes in vacuolar morphology. Consistent with their both having NPC assembly defects, *nup116ΔGLFG* cells exhibited synthetic growth defects when combined with *nup120Δ* or *nup133Δ* mutants (Fig. 2 C).

**Figure 2:**
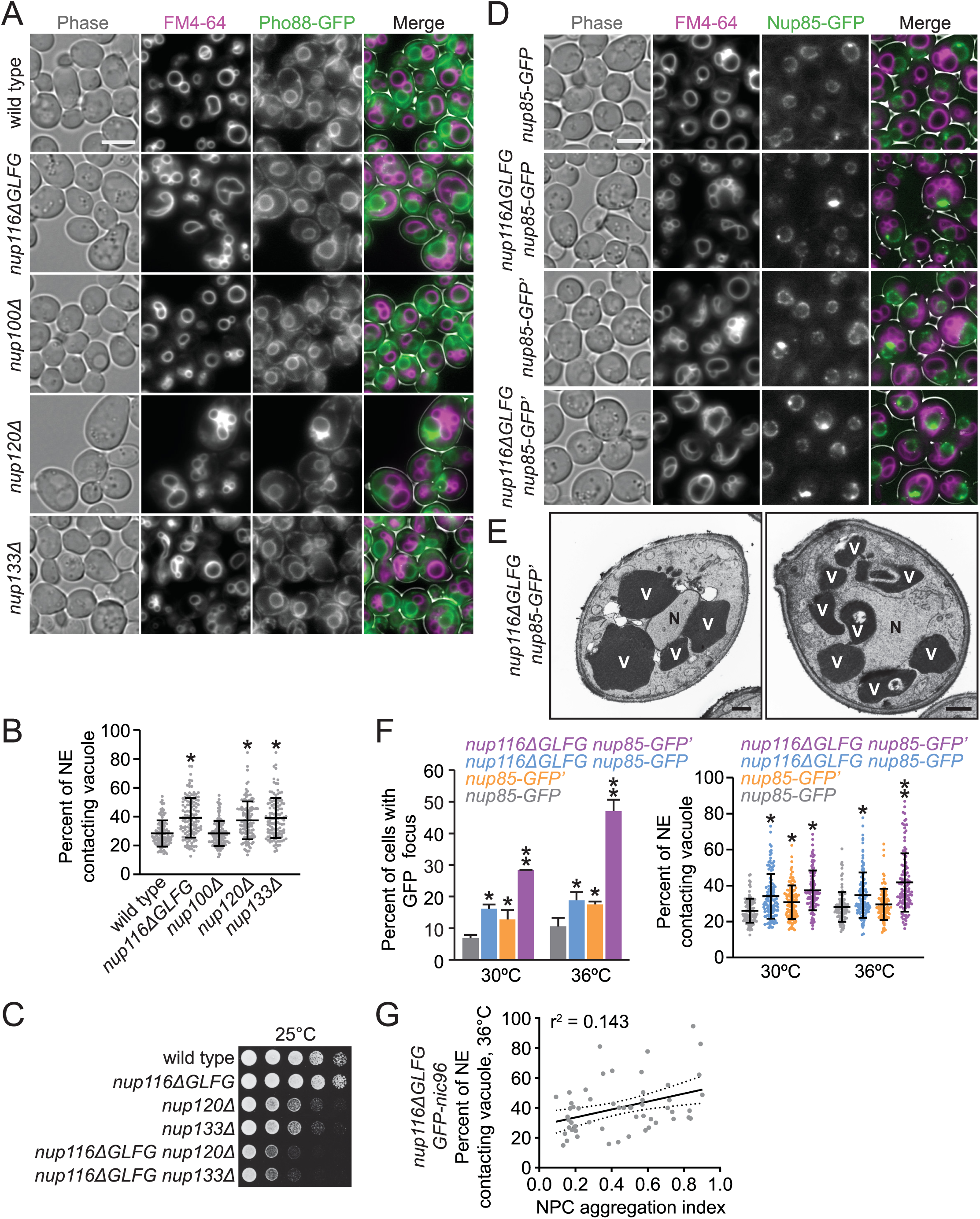
Several nucleoporin mutations that disrupt NPC assembly also increase NE-vacuole contacts. **(A)** FM4-64-stained *pho88-GFP* strains grown at 30°C. Bar, 5 μm. **(B)** NE-vacuole contacts at 30°C. Each dot represents contacts from one cell; 40 cells per strain were analyzed from 3 separate experiments. Asterisk indicates a p-value of less than 0.001 when compared to wild type using Dunn’s post-hoc test. **(C)** Serial dilutions of cells onto YPD plates. **(D)** FM4-64 labeled *nup85-GFP* or *nup85-GFP*’ (refer to Methods for details regarding strain construction) cells grown at 36°C. Bar, 5 μm. **(E)** TEM images of *nup116ΔGLFG nup85-GFP’* strains grown at 36°C. Nuclei are labeled with a black “N”, while vacuoles are labeled with a white “V”. Bars, 500 nm. **(F) (Left)** Percentage of cells with at least one GFP focus at 30 or 36°C. One asterisk indicates a p-value of less than 0.05 when compared to *nup85-GFP*, two asterisks indicate a p-value of less than 0.05 when compared to all other values using an unpaired two-tailed Student’s t-test for a particular temperature with an n of at least 100 cells from 3 independent experiments. **(Right)** NE-vacuole contacts from strains grown at 30 or 36°C. Each dot represents one cell with 40 cells per strain analyzed from 3 different experiments. One asterisk indicates a p-value of less than 0.01 when compared to *nup85-GFP*, two asterisks indicate a p-value of less than 0.05 when compared to all other values using Dunn’s post-hoc test for a particular temperature. **(G)** NE-vacuole contacts vs. NPC aggregation index for *GFP-nic96 nup116ΔGLFG* cells grown at 36°C. Each dot represents values from one cell with a total of 50 analyzed. Solid line represents the linear regression of the data, while dotted lines represent 95% confidence interval.

To confirm Nup116-dependent NPC clustering and NE-vacuole interactions using a different NPC marker than GFP-Nic96, *nup85-GFP’* from the yeast GFP collection (Huh et al., 2003), (arbitrarily and henceforth referred to as *nup85-GFP’*), was introduced into *nup116ΔGLFG* cells (Fig. 2, D and E; Fig. S3 A). Surprisingly, greater amounts of NE-vacuole contacts were detectable in *nup116ΔGLFG nup85-GFP’* cells compared to *nup116ΔGLFG GFP-nic96* cells at 36°C, with some *nup116ΔGLFG nup85-GFP’* cells having nuclei that were nearly completely surrounded by vacuoles (Fig. 2, D-F). Even *nup85-GFP’* cells exhibited somewhat increased NE-vacuole contacts, indicating that altering the function of *NUP85* with this particular GFP epitope impacts vacuole morphology; *nup85-GFP’* was at least partially compromised given its modest growth defect at 38.5°C and its synthetic growth defect with *nup116ΔGLFG* mutants (Fig. S1 D).

As a comparison, the pYM25 (Janke et al., 2004) vector was also used to generate *nup85-GFP* and *nup116ΔGLFG nup85-GFP* strains with a shorter linker between the *nup* and *GFP* and with yeast codon-optimized GFP sequence. At both 30 and 36°C, *nup116ΔGLFG nup85-GFP* mutants exhibited less severe NPC clustering relative to *nup116ΔGLFG nup85-GFP’* mutants, and fewer NE-vacuole interactions at 36°C (Fig. 2, D-F). DNA sequencing revealed no differences in the *NUP85* coding region between *nup85-GFP* and *nup85-GFP’*, suggesting the linker and/or GFP sequence of *nup85-GFP* were responsible for promoting proper function. We concluded that partially inhibiting the function of *NUP85* can impact NPC clustering as well as NE-vacuole interactions, and increasing the severity of NPC clustering tends to cause greater changes in vacuole morphology.

We did note, however, that the exact level of NPC clustering did not completely correlate with increased NE-vacuole contacts. For example, *nup85-GFP’* and *nup116ΔGLFG nup85-GFP* strains both exhibited roughly similar proportions of the cell population with GFP foci, but NE-vacuole contacts were more abundant in *nup116ΔGLFG nup85-GFP* cells at 36°C (Fig. 2 F). Additionally, when comparing the severity of NPC clustering defects using an aggregation index (Casey et al., 2015; Niepel et al., 2013) (higher index indicates more severe clustering) with the extent of NE-vacuole interactions in *nup116ΔGLFG GFP-nic96* cells at the single cell level (Fig. 2 G), there was no obvious correlation between these two variables. Thus, increased NE-vacuole contacts were observed in several *nup* mutations that coincide with NPC assembly defects and NE herniations; though at the single-cell level, the presence and severity of GFP-Nup clustering did not necessarily correlate with the extent of NE-vacuole interactions.

### NVJ factors and LDs are enriched at NE-vacuole contacts in NPC assembly mutants

We hypothesized that the extensive NE-vacuole contacts in *nup116ΔGLFG* mutants necessitated changes in vacuole membrane composition that could be reversed using chlorpromazine, which increases membrane fluidity and tends to enhance vacuole fusion in vitro (Fratti et al., 2007). Following a 5 min chlorpromazine treatment, *GFP-nic96* and *GFP-nic96 nup116ΔGLFG* vacuoles tended to form one large sphere that minimally interacted with the NE (Fig. 3 A), unlike chlorpromazine-insensitive *GFP-nic96 ypt7Δ* mutants. Despite these physical changes to the vacuole and consistent with their increased association under normal conditions, however, we observed some *GFP-nic96 nup116ΔGLFG* cells that still retained FM4-64-positive membranes around the NE (arrows, Fig. 3 A). This suggested that NE-associated factors can still at least transiently interact with vacuole lipids after treatment.

**Figure 3:**
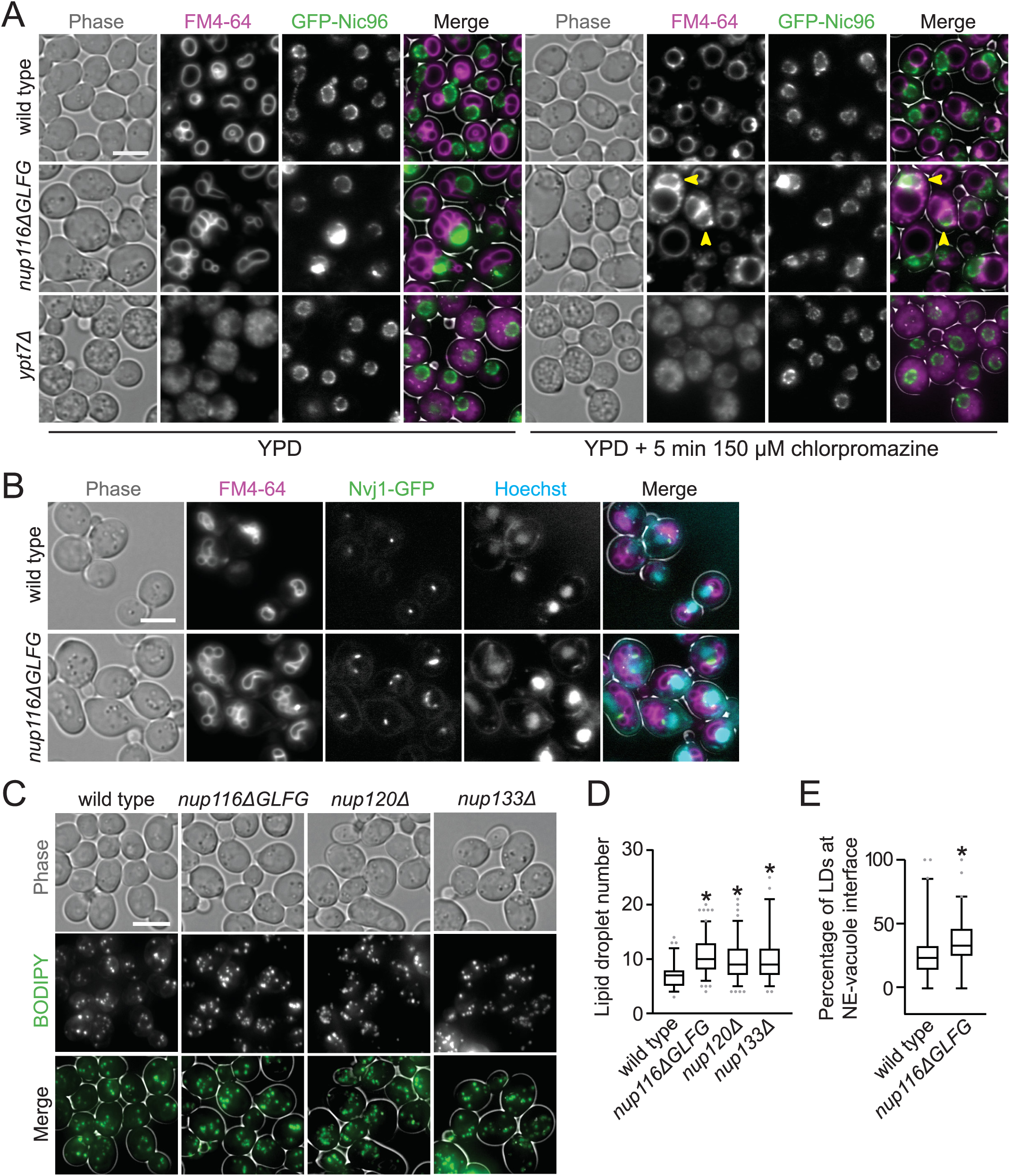
Compositional changes to NE-vacuole interfaces and increased lipid droplet levels in NPC assembly mutants. **(A)** FM4-64-stained *GFP-nic96* yeast grown at 36°C were imaged before (left) and after (right) incubation with 150 µm chlorpromazine for 5 min. Arrows point to cells that still have significant FM4-64 signal around NE following treatment. Bar, 5 µm. **(B)** FM4-64- and Hoechst-stained *nvj1-GFP* yeast grown at 36°C. Bar, 5 µm. **(C)** Max project images of BODIPY-stained yeast grown at 30°C. Bar, 5 µm. **(D)** Total LD number in the listed strains grown at 30°C. Asterisk indicates a p-value of less than 0.0001 when comparing to wild type using Dunn’s post-hoc test. Box and whiskers represent 2.5-97.5 percentile. **(E)** Percentage of LDs that are at NE-vacuole interface in the listed strains at 34°C. Asterisk indicates a p-value of less than 0.001 when compared to wild type using a two-tailed Mann-Whitney U test. Box and whiskers represent 2.5-97.5 percentile.

The interorganelle contacts present in *nup116ΔGLFG* mutants were suggestive of NVJs. To test this, the localization of Nvj1-GFP (Pan et al., 2000) and Mdm1-GFP (Henne et al., 2015) were analyzed to determine whether protein composition at these interfaces was also altered. Unlike wild type cells, which generally displayed a single focus of Nvj1-GFP at NE-vacuole interfaces, *nup116ΔGLFG* mutants often exhibited extensive regions of Nvj1-GFP at NE-vacuole contacts at 36°C (Fig. 3 B). Similar results were observed with Mdm1-GFP at 30°C, though Mdm1-GFP was only barely detectable in wild type and *nup116ΔGLFG* cells (Fig. S3 B) due to its low expression level (Ghaemmaghami et al., 2003). Because NVJs can serve as sites of LD biogenesis (Hariri et al., 2018) and increased LD levels have been previously observed in NPC assembly mutants (Hodge et al., 2010), total amounts of LDs were measured using BODIPY in *nup116ΔGLFG*, *nup120Δ*, and *nup133Δ* mutants. LD number was significantly increased in these mutant strains (Fig. 3, C and D), and LDs were enriched at NE-vacuole contact sites (Fig. 3 E; Fig. S3 C). These results indicated that NVJs expand and LD number increases when NPC assembly is compromised, producing significant compositional changes to the NE that promote vacuole interactions.

### NVJs and LDs cooperatively promote viability and NPC formation in assembly mutants

We reasoned that the increased NE-vacuole contacts and LDs in specific *nup* mutants could either be a deleterious/neutral effect caused by altered NPC function, or a secondary protective response that mitigates complications that arise from disrupted NPC assembly. To investigate these competing hypotheses, *NVJ1* and *MDM1* were deleted in *nup116ΔGLFG GFP-nic96* cells, as were *LRO1* and *DGA1*, two redundant acetyltransferase genes required for LD formation (Oelkers et al., 2002; Petschnigg et al., 2009). Synthetic growth defects in quadruple mutants would be consistent with a model wherein LDs or NVJs ameliorate NPC assembly stress, whereas other growth phenotypes would suggest NVJs or LDs are neutral/harmful effects. Deletion of *NVJ1* and *MDM1* impaired NE-vacuole contacts in *GFP-nic96* and *GFP-nic96 nup116ΔGLFG* cells (Fig. 4 A), and *DGA1*- and *LRO1*-deficient strains exhibited reduced LD numbers (Fig. S4 A).

**Figure 4:**
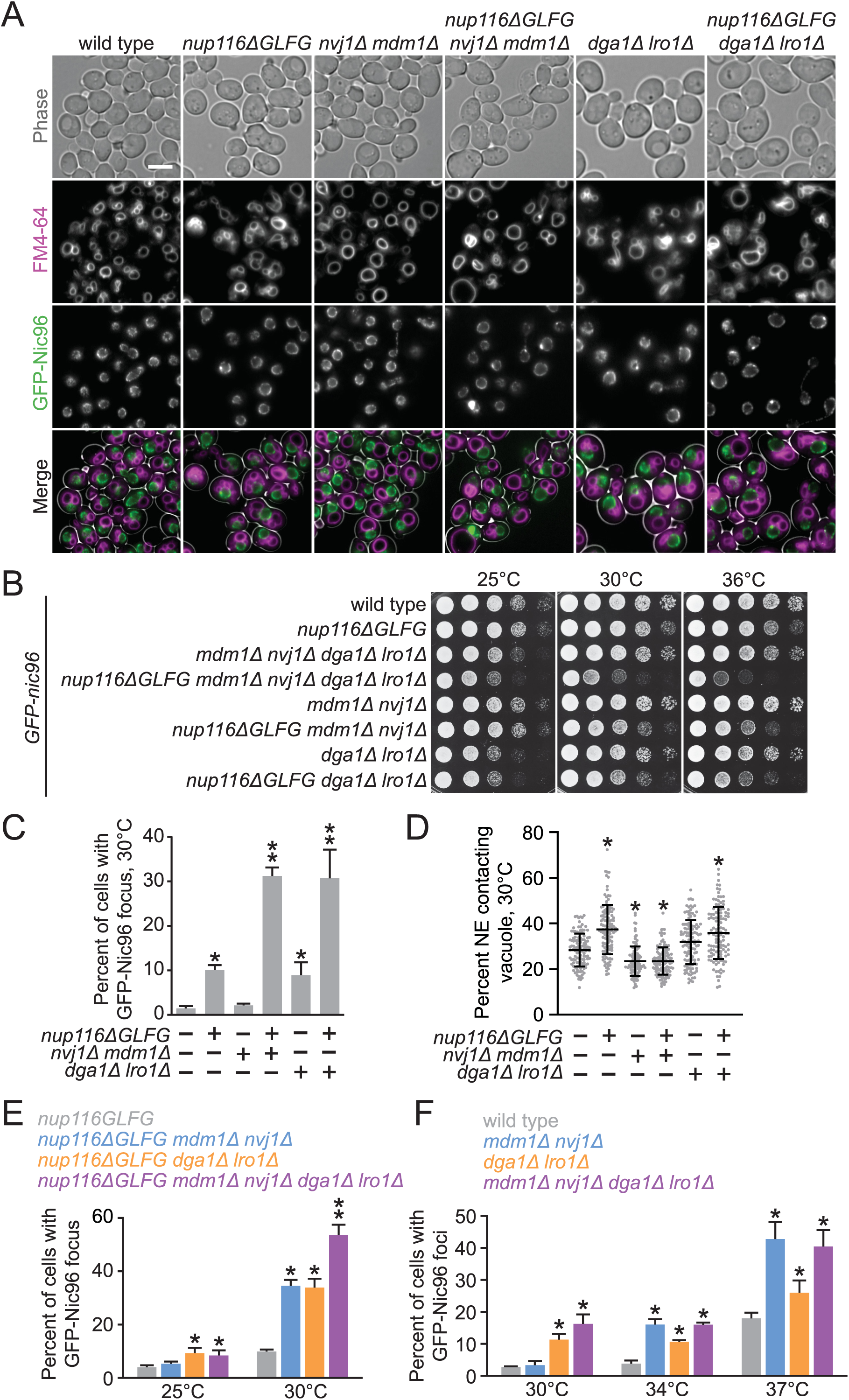
NVJs and LDs promote NPC formation and proper growth in NPC assembly mutants. **(A)** FM4-64-stained *GFP-nic96* strains grown at 30°C. Bar, 5 μm. **(B)** Serial dilutions of *GFP-nic96* yeast strains on YPD plates. **(C)** Percentage of cells grown at 30°C that contain at least one GFP-Nic96 focus. Asterisk indicates a p-value of less than 0.005 when compared to wild type using an unpaired two-tailed Student’s t-test with an n of at least 100 cells from 3 independent experiments; two asterisks indicate a p-value of less than 0.03 when also compared to *nup116ΔGLFG* cells. **(D)** NE-vacuole interactions from *GFP-nic96* strains grown at 30°C. Each dot represents value from one cell; 40 cells from 3 independent experiments were analyzed. Asterisk indicates a p-value of less than 0.01 when compared to wild type using Dunn’s post-hoc test. **(E)** Percentage of cells grown at 25 or 30°C that contain at least one GFP-Nic96 focus. One asterisk indicates a p-value of less than 0.05 when compared to *nup116ΔGLFG* cells at the same temperature using an unpaired two-tailed Student’s t-test with an n of at least 100 cells from 3 independent experiments; two asterisks indicate a p-value of less than 0.05 when also compared to *nup116ΔGLFG mdm1Δ nvj1Δ* and *nup116ΔGLFG dga1Δ lro1Δ* cells. **(F)** Percentage of cells grown at 30, 34, or 37°C that contain at least one GFP-Nic96 focus. One asterisk indicates a p-value of less than 0.05 when compared to wild type cells at the same temperature using a two-tailed Student’s t-test with an n of at least 100 cells from 3 independent experiments; *mdm1Δ nvj1Δ dga1Δ lro1Δ* mutants do not exhibit p-values of less than 0.05 when compared to other strains.

Both *GFP-nic96 nup116ΔGLFG nvj1Δ mdm1Δ* and *GFP-nic96 nup116ΔGLFG dga1Δ lro1Δ* mutants displayed substantial growth defects relative to *GFP-nic96 nup116ΔGLFG* cells at 34 and 36°C (Fig. 4 B). In contrast, growth of *GFP-nic96 nvj1Δ mdm1Δ* mutants was indistinguishable from wild type at all temperatures tested, while *GFP-nic96 dga1Δ lro1Δ* cells had growth defects at 25°C. Interestingly, LD- and NVJ-deficient *GFP-nic96* cells had only minor growth phenotypes at higher temperatures, but produced additive growth defects in *GFP-nic96 nup116ΔGLFG* mutants in a *NUP116*-dependent manner (Fig. S4 B), suggesting NVJs and LDs regulate viability at least partially independent of one another. Similar synthetic growth defects were observed in *nup133Δ mdm1Δ*, *nup133Δ nvj1Δ* and *nup133Δ dga1Δ lro1Δ* mutants (Fig. S4 C; note-*nup133Δ mdm1Δ nvj1Δ* mutants could not be successfully isolated). Thus, NVJs and LDs were required for maximal viability in several *nup* mutants with disrupted NPC assembly, consistent with a model wherein LDs and NE-vacuole contacts cooperate to ameliorate NPC assembly stress.

We next tested whether NPC assembly was disrupted in NVJ- or LD-deficient *GFP-nic96 nup116ΔGLFG* mutants by measuring foci formation and NE-vacuole contacts at 30°C, a temperature that allowed detection of subtle changes in NPC assembly. While deletion of *MDM1* and *NVJ1* did not impact NPC formation in *GFP-nic96* cells despite decreasing NE-vacuole interactions, 31% of *GFP-nic96 nup116ΔGLFG mdm1Δ nvj1Δ* mutants displayed GFP-Nic96 foci, a three-fold increase compared to *GFP-nic96 nup116ΔGLFG* cells (Fig. 4, A-D). 9% of *GFP-nic96 dga1Δ lro1Δ* mutants exhibited NPC assembly defects, and potentially because of this or to compensate for their decreased LD content, may also have increased NE-vacuole contacts (p<0.05 when using a Mann-Whitney U test, but not Dunn’s post-hoc test, when compared to wild type); 30% of *GFP-nic96 nup116ΔGLFG dga1Δ lro1Δ* mutants had GFP-Nic96 foci. Additionally, NVJ-deficient *nup116ΔGLFG* mutants tended to exhibit more NE herniations (Fig. S4 D). Overall, NE-vacuole contacts promoted NPC assembly and proper NE morphology in *nup116ΔGLFG* cells despite having no discernable impact in wild type cells at 30°C. Inhibiting LD formation did impair NPC formation in otherwise wild type cells, but produced synthetic assembly defects in *nup116ΔGLFG* mutants. Disrupting NVJs did not significantly impact total numbers of LDs, although fewer were present at the nuclear rim (Fig. S4 E).

To test how NVJs and LDs regulate NPC assembly together, foci formation was assessed in *GFP-nic96 nup116ΔGLFG mdm1Δ nvj1Δ dga1Δ lro1Δ* and *GFP-nic96 mdm1Δ nvj1Δ dga1Δ lro1Δ* strains at several temperatures. While NVJ- and LD-deficient *nup116ΔGLFG* mutants did not have additive NPC clustering defects at 25°C, nearly 55% of the mutant cells displayed GFP-Nic96 foci at 30°C compared to ∼30% of *nup116ΔGLFG* cells deficient for LDs or NVJs (Fig. 4 E), consistent with their additive growth defects (Fig. 4 B). Thus, LDs and NVJs separately inhibited GFP-Nic96 foci formation in the context of perturbed NPC assembly. Interestingly, NVJs and LDs did not appear to function in a partially redundant manner in wild type cells, as *GFP-nic96 mdm1Δ nvj1Δ dga1Δ lro1Δ* mutants did not exhibit clustering defects that are statistically more severe than *GFP-nic96 mdm1Δ nvj1Δ* or *GFP-nic96 dga1Δ lro1Δ* mutants at all temperatures tested (Fig. 4 F). We speculated that the NPC assembly defects at 34 and 37°C in *GFP-nic96 mdm1Δ nvj1Δ* cells result from mild assembly stress caused by elevated temperature and/or GFP-tagging *NIC96*, though NVJs could simply be more critical to regulating NPC assembly in wild type cells at higher temperatures.

### NVJs and autophagy factors mediate *PEP4*-dependent NE remodeling in *nup116ΔGLFG* mutants

Nitrogen starvation causes PMN to occur following expansion of NVJs (Kvam and Goldfarb, 2004). We therefore hypothesized that regions of the NE could be undergoing vacuole-mediated remodeling/degradation in NPC assembly mutants. Anti-GFP immunoblots of lysates derived from Nup85-GFP, GFP-Nic96, or Pho88-GFP strains were analyzed to examine whether degradation of any of these proteins was *NUP116*-dependent (Fig. 5 A). Pho88-GFP was used as a control; if only Nups were degraded, we assumed this would likely indicate Nup degradation occurs prior to insertion into the NE and would be caused by disrupted NPC assembly as opposed to NE remodeling.

**Figure 5:**
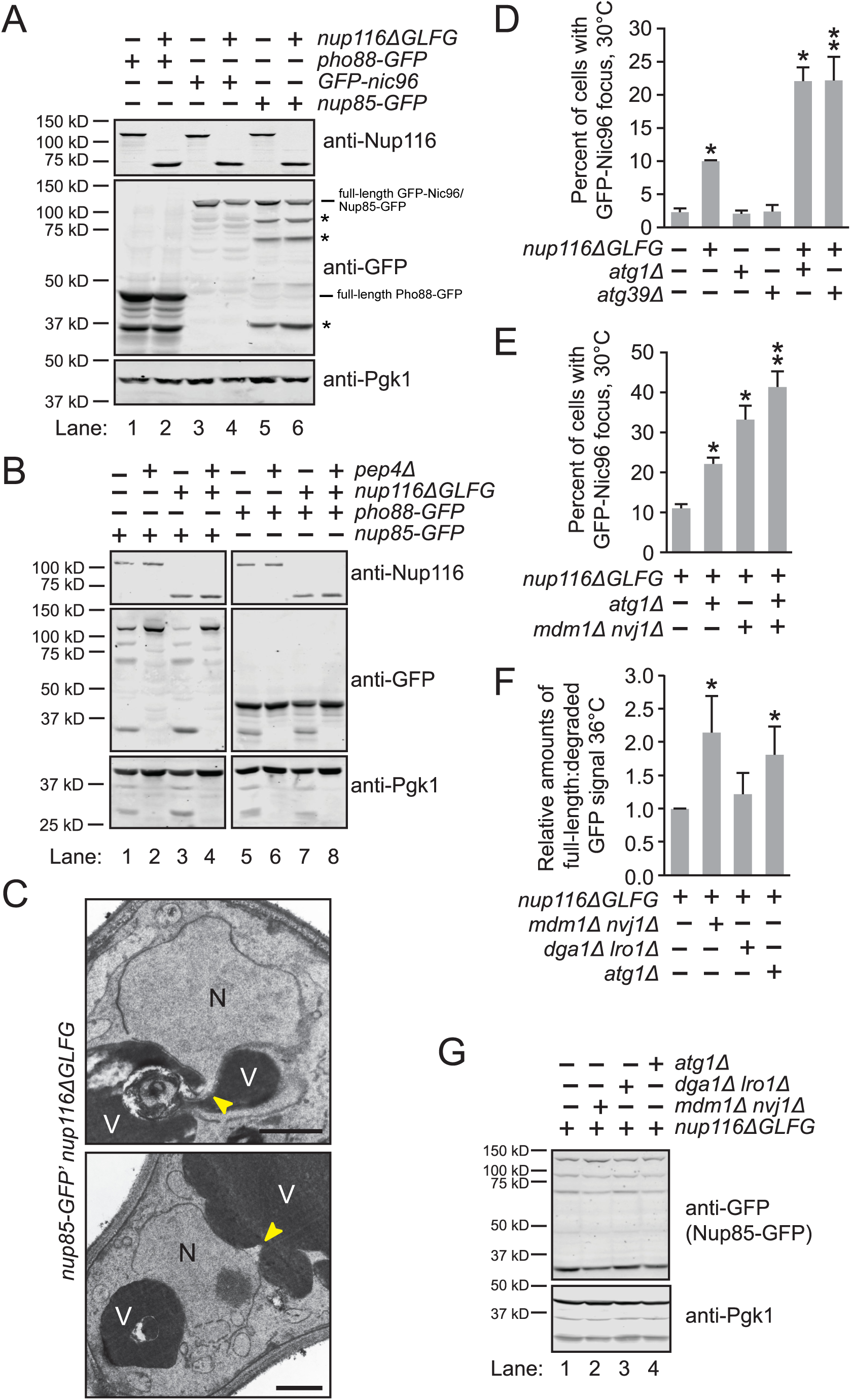
NE-vacuole interactions and autophagy factors promote NE remodeling in *nup116ΔGLFG* mutants. **(A)** and **(B)** Western blots of lysates from the listed strains. Pgk1 is a cytoplasmic protein that was used as a loading control. Asterisks indicate major degradation products. **(C)** TEM images of *nup85-GFP’ nup116ΔGLFG* cells grown at 36°C. Nuclei are labeled with a black “N” and vacuoles are labeled with a white “V”. Potential sites of NE remodeling are labeled with a yellow arrow. Bars, 500 nm. **(D)** Percentage of cells grown at 30°C with at least one GFP-Nic96 focus. One asterisk indicates a p-value of less than 0.01 when compared to wild type using an unpaired two-tailed Student’s t-test with an n of at least 100 cells from 3 independent experiments; two asterisks indicate a p-value of less than 0.05 when also compared to single mutants. **(E)** Percent of cells from different *GFP-nic96 nup116ΔGLFG* strains with at least one GFP-Nic96 focus. One asterisk indicates a p-value of less than 0.001 when compared to *nup116ΔGLFG* cells using an unpaired two-tailed Student’s t-test with an n of at least 100 cells from 5 independent experiments, while two asterisks indicate a p-value of less than 0.01 when compared to all other strains. **(F)** Quantification of relative degradation levels of Nup85-GFP in the listed *nup116ΔGLFG* strains at 36°C. Asterisks indicate a p-value of less than 0.02 when compared to *nup116ΔGLFG* cells using an unpaired two-tailed Student’s t-test with an n of 5. **(G)** Western blots of *nup85-GFP nup116ΔGLFG* lysates from cells grown at 36°C; this data and other independent experiments were used to generate data in F. Asterisks indicate major degradation products.

Interestingly, the total levels of all three proteins were decreased in *nup116ΔGLFG* lysates, and for C-terminally tagged Nup85-GFP and Pho88-GFP, substantially more degradation products were detectable in *nup116ΔGLFG* lysates (Fig. 5 A, compare lanes 1 and 2, 3 and 4, 5 and 6). Degradation of Nup85-GFP and Pho88-GFP required the vacuolar peptidase *PEP4* (Fig. 5 B), suggesting processing occurs in vacuoles. Unlike starvation-induced PMN, we were unable to detect large blebs of nuclei in vacuoles, though smaller regions of the NE were infrequently observed that potentially were directly remodeled near vacuoles (Fig. 5 C). Based on this, we concluded that NPC assembly mutants undergo vacuole-dependent NE remodeling that is functionally distinct from PMN.

The kinase Atg1 and several other factors are required for starvation-induced PMN that occurs at NVJs (Krick et al., 2008), while the autophagy receptor Atg39 is necessary for rapamycin-induced nucleophagy (Mochida et al., 2015). It is possible that these autophagy proteins could be mediating NE remodeling to prevent NPC clustering in *nup116ΔGLFG* mutants. Like NVJs and LDs, we predicted significantly more GFP-Nic96 foci would be observed in combinatorial mutants if *ATG1* and/or *ATG39* was necessary to prevent formation of NPC clusters. Analysis of GFP-Nic96 foci formation revealed that while neither *ATG1* nor *ATG39* significantly impacted NPC assembly, deletion of either gene along with Nup116’s GLFG domain doubled the levels of GFP-Nic96 foci at 30°C (Fig. 5 D; Fig. S5 A). Interestingly, these autophagy factors likely function independent of NVJs because *GFP-nic96 nup116ΔGLFG mdm1Δ nvj1Δ atg1Δ* mutants exhibited additive NPC clustering defects compared to *GFP-nic96 nup116ΔGLFG mdm1Δ nvj1Δ* cells (Fig. 5 E). Additionally, NE-vacuole interactions in *nup116ΔGLFG* mutants were unaffected by deletion of *ATG1* or *ATG39* (Fig. S5 B). Unlike NVJ and LD genes, *GFP-nic96 nup116ΔGLFG atg1Δ* and *GFP-nic96 nup116ΔGLFG atg39Δ* mutants did not display additive growth defects (Fig. S5 C), indicating that the extent of NPC clustering does not necessarily correlate with growth rate. Thus, *ATG1* and *ATG39* were necessary to limit NPC clustering in *nup116ΔGLFG* mutants along with NVJs, and neither gene impacted the viability of NPC assembly mutants under normal growth conditions.

To directly assess whether autophagy, NE-vacuole contacts, and LDs were required for NE remodeling in *nup116ΔGLFG* cells, Nup85-GFP degradation was compared in strains lacking *MDM1* and *NVJ1*, *DGA1* and *LRO1*, or *ATG1* at 36°C. Significantly less Nup85-GFP degradation occurred in *nup85-GFP nup116ΔGLFG mdm1Δ nvj1Δ* and *nup85-GFP nup116ΔGLFG atg1Δ* lysates compared to *nup85-GFP nup116ΔGLFG* lysates, though degradation was unchanged in *nup85-GFP nup116ΔGLFG dga1Δ lro1Δ* lysates (Fig. 5, F and G; compare lanes 1-4). These results indicated that NVJ and autophagy factors, but not LDs, are required for degradation of NE components in *nup116ΔGLFG* vacuoles.

We speculated that NVJ-mediated remodeling could be occurring in *nup116ΔGLFG* cells at a relatively constant level that prevents formation and/or expansion of NPC clusters, or in a manner that promotes direct removal of clustered NPCs. To distinguish between these possibilities, *GFP-nic96 nup116ΔGLFG* cells were monitored for several hours using a microfluidic chamber to determine if GFP-Nic96 foci were ever observed being cleared from the NE (Video 1). Since we did not observe the removal of any obvious NPC clusters over 4.5 hours, we concluded that *NVJ1*- and *MDM1*-mediated NE-vacuole contacts continuously remodel the NE under conditions of NPC assembly stress to prevent NPC cluster formation/expansion.

### *CHM7* differentially affects *nup116ΔGLFG* and *vps4Δ* mutants

Similar to SINC NPC clusters, which form when a *VPS4*-mediated NPC surveillance pathway is inhibited (Webster et al., 2014), GFP-Nic96 foci were not present in every mutant *nup116ΔGLFG* cell (Fig. 1 B). We therefore tested whether *nup116ΔGLFG* foci are retained in mother yeast cells like SINC NPCs. Comparison of NPC aggregation in mother vs. daughter *GFP-nic96 nup116ΔGLFG* nuclei in cells that had not yet fully divided and contained at least one focus at 34°C revealed that NPC aggregation indices were greater in mother cells (Fig. 6, A and B). Consistent with this data, NPC aggregation increased as *nup116ΔGLFG* mutants replicatively aged at 34°C relative to wild type cells (Fig. 6 C). Importantly, these experiments indicated that NPC aggregates in *nup116ΔGLFG* cells tend to be retained in mother cells.

**Figure 6:**
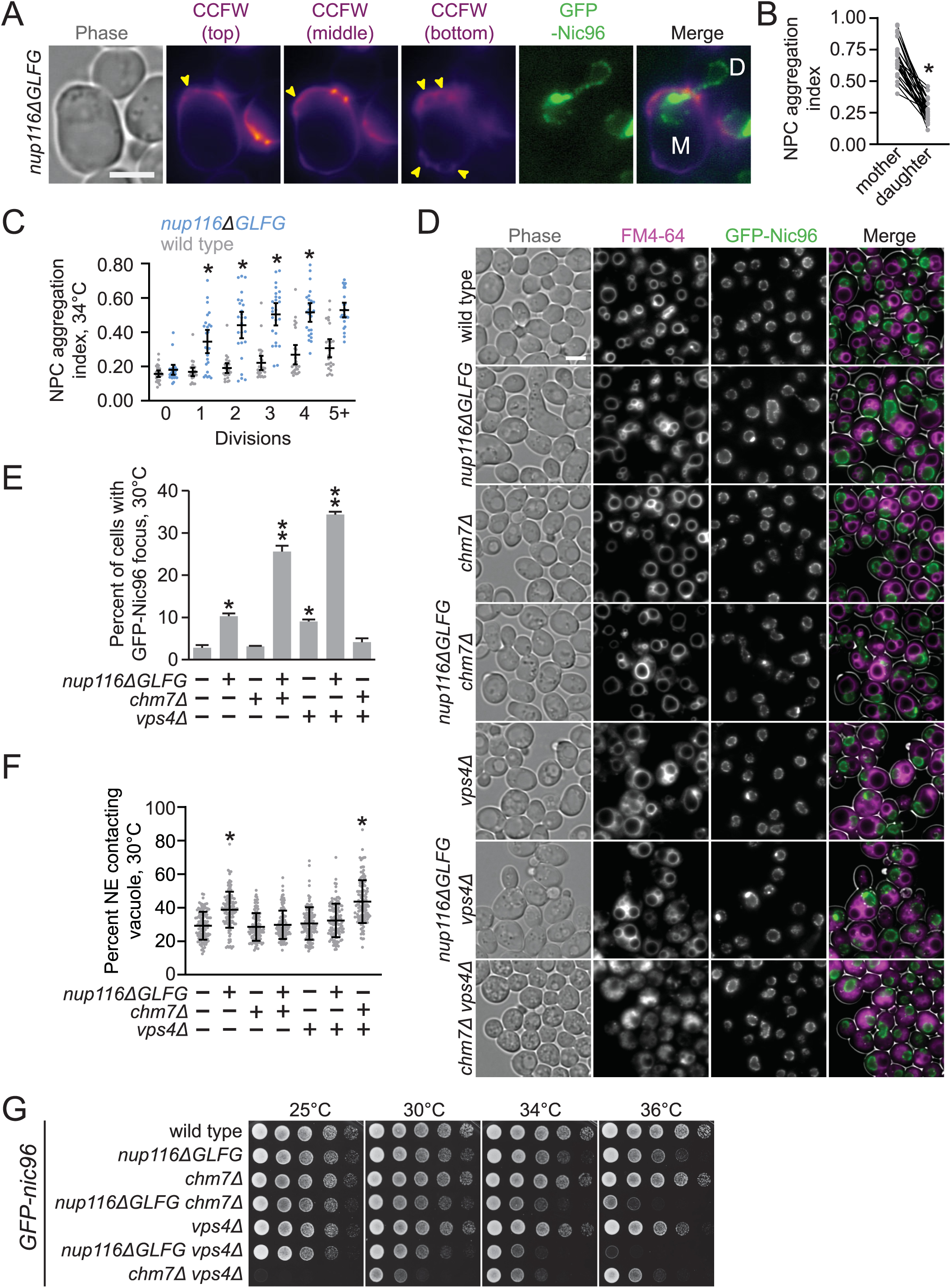
NPC clustering in *nup116ΔGLFG* cells is *CHM7*-independent. **(A)** Calcofluor white (CCFW)- stained *GFP-nic96 nup116ΔGLFG* cell grown to at 34°C that underwent 6 divisions. Mother cell is labeled with an “M”, while undivided daughter is labeled with a “D”; arrows point to bud scars. Scale bar, 5 µm. **(B)** NPC aggregation indices from undivided mother cells with mother and daughter nuclei. Each left dot represents one mother nucleus, connected with its corresponding daughter nucleus on the right. Asterisk indicates a p-value of less than 0.001 when compared using a paired two-tailed Wilcoxon test. **(C)** NPC aggregation index vs division number for *GFP-nic96* strains at 34°C. Each dot represents values from one cell; asterisk indicates a p-value of less than 0.05 when compared to wild type using Dunn’s post-hoc test for given division number. **(D)** FM4-64-stained *GFP-nic96* cells grown at 30°C. Bar, 5 µm. **(E)** Percentage of cells with at least one GFP-Nic96 focus from cells grown at 30°C. One asterisk indicates a p-value of less than 0.01 when compared to wild type using an unpaired two-tailed Student’s t-test with an n of at least 100 cells from 3 experiments; two asterisks indicate a p-value of less than 0.01 when also compared to *nup116ΔGLFG* or *vps4Δ* cells. **(F)** NE-vacuole contacts in the listed *GFP-nic96* strains grown at 30°C. Each dot corresponds to the value from an individual cell. Asterisk indicates a p-value of less than 0.01 when compared to wild type using Dunn’s post-hoc test with an n of 40 cells from 3 experiments. **(G)** Serial dilutions of *GFP-nic96* strains on YPD plates.

Formation of SINC foci in *vps4Δ* mutants is abolished in the absence of the ESCRT subunit Chm7 (Webster et al., 2016), likely because clustering is promoted by improper compartmentalization of Chm7 (Thaller et al., 2019). GFP-Nic96 foci and NE-vacuole contacts were measured in *GFP-nic96 nup116ΔGLFG chm7Δ* cells at 30°C to test whether foci formation in *nup116ΔGLFG* mutants also required *CHM7*. Surprisingly, instead of abrogating GFP-Nic96 foci formation as it does in *vps4Δ* cells (Fig. 6, D and E), deletion of *CHM7* increased GFP-Nic96 foci formation ∼150% in *nup116ΔGLFG* cells without impacting NPC assembly in otherwise wild type cells. This correlated with a significant loss of NE-vacuole contacts in *GFP-nic96 nup116ΔGLFG chm7Δ* mutants (Fig. 6 F); similar effects were observed when *VPS4* was deleted in *GFP-nic96 nup116ΔGLFG* mutants. The loss of NE-vacuole interactions by deletion of *CHM7* or *VPS4* suggested that Chm7 and Vps4 are somehow required to alter NE/vacuole composition so that these interorganelle interactions can occur in *nup116ΔGLFG* mutants. Consistent with a loss of NE-vacuole interactions, *GFP-nic96 nup116ΔGLFG chm7Δ* and *GFP-nic96 nup116ΔGLFG vps4Δ* cells exhibited significant synthetic growth defects (Fig. 6 G). In sum, NPC clusters in *nup116ΔGLFG* mutants formed in a *CHM7*-independent manner distinct from those observed in *vps4Δ* mutants, suggesting NVJ-mediated regulation of NPC assembly functions in a separate pathway than *VPS4*-mediated assembly surveillance.

## Discussion

Using *Saccharomyces cerevisiae* mutants that induce NPC assembly defects as models, we demonstrate that interorganelle contacts between the NE and vacuoles expand in response to NPC assembly stress. These contact sites are enriched with LDs as well as proteins that promote the formation of NVJs. We also show that NE-vacuole contacts cooperate with LDs to maintain viability and stabilize NPC formation when assembly is perturbed. Finally, autophagy factors function with NVJs to remodel the NE in NPC assembly mutants. Importantly, this work reveals that interorganelle contacts between the NE and vacuole perform novel functions to ameliorate NPC assembly stress.

Results presented herein suggest a model whereby NPC assembly is compromised in *nup116ΔGLFG* mutants, particularly at elevated temperatures, leading to expansion of NVJs that cooperate with LDs to maintain cell viability as well as remodel the NE in a *PEP4*-dependent manner with the autophagy factors *ATG1* and *ATG39* (Fig 7). We propose that both of these NVJ-dependent processes are necessary to promote proper NPC assembly. Although NE-vacuole contacts do not impact NPC assembly in wild type cells at 30°C, they play essential roles in mitigating stress induced by several *nup* mutants that disrupt NPC assembly (Fig. 4, Fig. S4) and potentially temperature-dependent assembly stress (Fig. 4 F).

**Figure 7:**
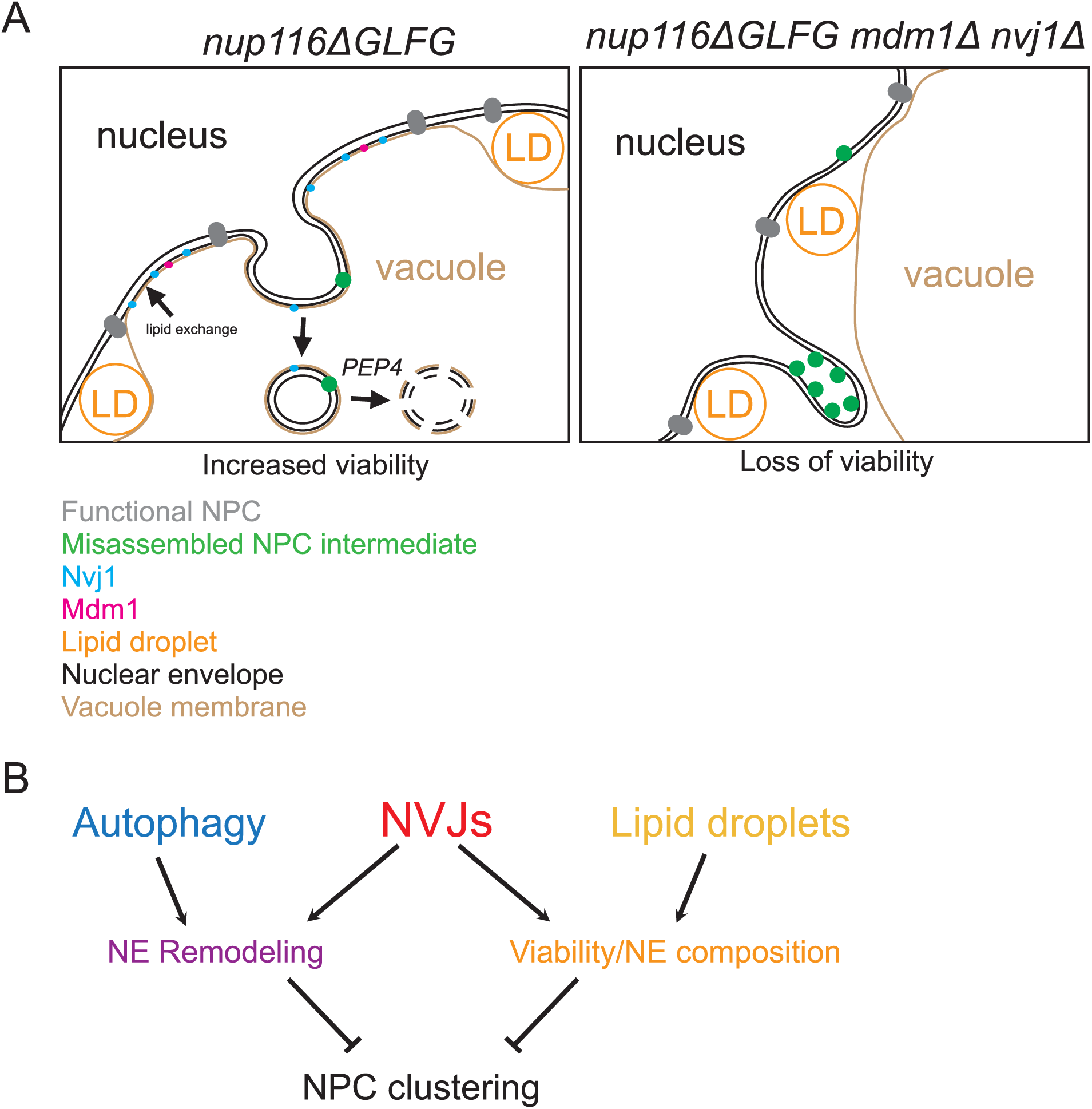
NE-vacuole interactions mediate dual responses to NPC assembly stress. **(A)** and **(B)** Cartoons depicting proposed functions of NVJs in *nup116ΔGLFG* cells. NVJs containing Nvj1 and Mdm1 expand around the NE in response to misassembled NPCs, and lipid droplets also are enriched at these contacts. NVJs and LDs promote viability together, potentially through lipid exchange that stabilizes the NE. Additionally, NVJs promote continuous vacuole-mediated remodeling of the NE, potentially degrading misassembled NPC intermediates that form in the inner NE. Separately, the autophagy factors *ATG1* and *ATG39* also mediate remodeling of the NE. Together, these dual functions of NVJs enhance proper NPC formation in mutants where NPC assembly is perturbed. In the absence of NVJs, more misassembled NPC intermediates accumulate in NE herniations that appear as clusters using fluorescence microscopy.

The observation that NE-vacuole contacts do not expand in *nup100Δ* mutants, which display increased NPC permeability (Lord et al., 2015; Timney et al., 2016; Popken et al., 2015) and tRNA export defects (Lord et al., 2017), suggests NVJ formation in *nup116ΔGLFG*, *nup120Δ*, *nup133Δ,* and *nup85-GFP’* mutants is related to perturbation of NPC assembly and not a general loss of NPC function. We also propose that vacuole membrane composition is significantly altered in NPC assembly mutants to accommodate the significant morphology changes necessary to maintain viability. In support of this possibility, we observe significantly more Nvj1-GFP and Mdm1-GFP at NE-vacuole interfaces in *nup116ΔGLFG* cells, as well as partial reversal of NVJ phenotypes in these mutants in response to chlorpromazine (Fig. 3).

We speculate that NVJs and LDs promote viability in *nup116ΔGLFG* and *nup133Δ* strains by serving as interfaces to promote lipid exchange that beneficially alters NE composition in response to perturbations in NPC assembly. Although NVJs have not been specifically shown to mediate lipid transfer between the vacuole and NE, interorganelle contact sites often serve as sites of lipid exchange (Kornmann et al., 2009; Elbaz-Alon et al., 2014) due to their close apposition and enrichment of lipid-exchanging factors. The ergosterol- and PI(4)P-binding protein Osh1 (Manik et al., 2017), which is found at NVJs and interacts with Nvj1 (Jeong et al., 2017; Kvam and Goldfarb, 2004; Levine and Munro, 2001), belongs to the oxysterol-binding homology protein family that can aid in lipid transfer between membranes (de Saint-Jean et al., 2011). Additionally, LDs are thought to promote lipid transfer between organelles, potentially through “lipid bridges” (Schuldiner and Bohnert, 2017). Since deletion of NVJs or LDs significantly increases NPC clustering in *nup116ΔGLFG* cells (Fig. 4), we hypothesize that preventing changes in lipid composition via NVJs or LDs further inhibits NPC formation, resulting in more misassembled clustered NPCs and NE herniations.

Our experiments further demonstrate that NVJs and autophagy play important roles in remodeling the NE when NPC assembly is impaired (Fig. 5), leading to removal of NE domains that are enriched in NPC components like Nup85, but also contain other NE/ER membrane proteins such as Pho88. Following their removal, these domains are eventually degraded in the vacuole in a *PEP4*-dependent manner (Fig. 5). Based on their close apposition and potential intermediates observed using TEM, we propose that NVJs can serve as sites to directly promote removal of NE domains that contain misassembled NPCs (Fig. 7); NPC clusters are then less likely to form by removal of these domains in a fairly continuous manner (Video 1).

While the autophagy factor *ATG1* also remodels the NE, additive NPC clustering defects in triple mutants imply that they function at least somewhat separately from NVJs in doing so. We speculate the autophagy receptor Atg39 (Mochida et al., 2015) recognizes misassembled NPC intermediates, which are then sorted into autophagosomes and then transported to the vacuole in a largely NVJ-independent manner. Phosphorylation (Laurell et al., 2011) and ubiquitylation (Niño et al., 2016) are just some the of post-translational modifications that can be added to specific Nups, which may allow autophagy machinery and NVJs to distinguish them from assembled and transport-competent NPCs. We cannot exclude the possibility that NVJs utilize other autophagy proteins when remodeling the NE, and fully defining the cellular machinery required to deform the NE along with NVJs to promote remodeling will be an important future endeavor.

Recent work demonstrated that NPCs are degraded via an autophagy- and vacuole-dependent pathway in response to nitrogen starvation that exposes an Atg8-binding motif in Nup159 (Lee et al., 2020). Degradation appears somewhat upregulated in *nup120Δ* and *nup133Δ* mutants under normal growth conditions, suggesting this pathway is also utilized when NPC assembly is disrupted. We speculate this Atg8-dependent pathway could work separately with NVJs, other autophagy factors, and Chm7/Heh1 (Webster et al., 2016) to function as redundant mechanisms to respond to NPC assembly stress. NVJs are likely only utilized to remodel the NE under rich growth conditions, as deletion of *NVJ1* did not significantly impact NPC degradation during nitrogen starvation (Lee et al., 2020). More work will be required to ascertain whether Atg8-dependent autophagy is also utilized in *nup116ΔGLFG* mutants, particularly at higher temperatures that correlate with disrupted NPC assembly.

We propose expansion of NE-vacuole contacts and increased LD content constitutes a general cellular response to mitigate several types of sustained NPC assembly stress. For example, TEM images of other NPC assembly mutants clearly demonstrate many contain significantly more LDs, and less obviously, show accompanying changes in vacuole morphology (Makio et al., 2009; Ryan et al., 2003). Additionally, when the temperature-sensitive *brr6-1* allele is inactivated for several hours, NPC assembly is compromised and LD number increases; genetic experiments indicate these LDs are necessary for full viability of the mutant (Hodge et al., 2010). Short-term inactivation of Brr6, however, inhibits NPC assembly without altering lipid composition (Zhang et al., 2018), suggesting that LDs and potentially NVJs serve as a long-term cellular mechanism to compensate for perturbations in NPC formation. We do note that some mutants that produce NPC clusters, such as *vps4Δ* mutants, do not promote expansion of the vacuole along the NE. We speculate that this could be due to *VPS4*-dependent changes in the NE/vacuole membrane that prevent significant physical interactions with the NE. Further work will be required to resolve precisely if and how Vps4 and Chm7 modulate NVJs and other interorganelle contacts.

Deletion of Nup116’s GLFG domain in W303 cells prevents NPC biogenesis at 25°C when *NUP188* is inactivated, resulting in NE herniations without observable NPC clustering (Onischenko et al., 2017). The absence of clusters is likely caused by the complete abrogation of NPC synthesis, as prolonged inactivation of *NUP188* is lethal. Due to biochemical interactions of GLFG domains with several scaffolding Nups including Nup188, the failure of nup116ΔGLFG protein to incorporate into NPCs in the absence of *NUP188* (Onischenko et al., 2017), and that FG domains are inherently disordered (Hough et al., 2015; Denning et al., 2003), Nup116’s GLFG domain is proposed to directly facilitate NPC assembly by promoting aggregation of early NPC intermediates. We observe both NPC clusters and NE herniations in S288C *nup116ΔGLFG* cells that occur in a temperature-dependent manner, indicating deletion of Nup116’s GLFG domain produces significant NPC assembly defects without completely inhibiting NPC biogenesis. Although it seems likely that NPC assembly defects in S288C *nup116ΔGLFG* mutants at least partially result from disrupted Kap121-mediated transport, a direct role for its GLFG domain in facilitating NPC assembly via aggregation could also explain our results. Importantly, these models are not in any way mutually exclusive, and given the plethora of roles GLFG domains play in NPC function it may be difficult to define the precise causes of disrupted NPC biogenesis in *ΔGLFG* mutants.

NPC assembly and nuclear transport play essential roles in modulating yeast replicative life span (Lord et al., 2015, 2017; Rempel et al., 2019; King et al., 2019), and disrupted NPC function is associated with aging and neurological disorders in metazoans (D’Angelo et al., 2009; Toyama et al., 2013; Cruz and Cleveland, 2016). Given their impact on promoting proper NPC formation and viability in mutants with disrupted NPC assembly, it will be important to define how NE-vacuole interactions regulate the replicative aging process and potentially other stress conditions. Although metazoans have no obvious orthologous system to NVJs, their cells can produce nuclear LDs (Ohsaki et al., 2016) and regions of their nuclei can be degraded via autophagy (Dou et al., 2015), both of which could potentially be utilized to respond to perturbations in interphase NPC assembly. Importantly, our experiments establish novel and critical roles for NVJs in ameliorating NPC assembly defects, and provide mechanistic insights into how interorganelle contacts can be utilized to respond to stress/disease states.

## Materials and Methods

### Yeast strains and growth

A complete list of strains utilized for this study are listed in Supplementary Table 1. BY4741 and 4742 S288C strains were used for all experiments. Unless otherwise noted, yeast strains were grown in 1% yeast extract/2% peptone/2% dextrose (YPD) media. An S288C *nup116ΔGLFG* strain was generated by cloning the *nup116ΔGLFG* genome sequence from SWY2791 into SnaBI- and XhoI-digested pAG306-GPD-empty chr I (Addgene 41895) using Gibson Assembly master mix (NEB E2611L). The resulting vector was digested with SpeI (NEB R0133S) and transformed into BY4742 yeast. *URA3* positive transformants were then plated onto 5-FOA to excise full-length *NUP116*, which was confirmed using PCR and western blots. Other gene deletions were generated using PCR products derived from pFA6a-kanMX6 (Addgene 39296) or pFA6a-hphNT1 (Euroscarf P30347) vectors. *nup85-GFP*’ strains were derived from the GFP collection (Huh et al., 2003), itself derived from pFA6a-GFP(S65T)-His3MX6 (Addgene 41598), as were *nvj1-GFP* strains. pYM25 (Euroscarf P30237) was used to create *nup85-GFP* and *mdm1-GFP* strains. *GFP-nic96* strains were constructed by transforming strains with AflII-digested pSW950 (Bucci and Wente, 1998). Most combinatorial mutants were generated using mating and tetrad dissections.

### Plasmids

Other than the plasmids utilized to generate yeast strains described above, pRS315 (ATCC 77144) and pRS315*-NUP116* (Wente et al., 1992) were utilized for complementation.

### FM4-64 staining

1 ml of cells grown to mid-log phase in YPD were transferred to an amber 1.5 ml tube along with 1 µl of 8 mM FM4-64/DMSO for a final cell staining concentration of 8 µM. Cells were then stained at the temperature at which they grew overnight, along with shaking for 20-30 min. Following staining, cells were pelleted and washed two times with 1 ml fresh media (pre-warmed if necessary) and allowed to grow for an additional two hours prior to imaging, again at overnight growth temperature, unless otherwise noted for temperature-shift experiments.

### Fluorescence Microscopy

With the exception of Video 1, all images were acquired using a BX53 Olympus microscope equipped with a 100×/1.35 NA Olympus UPlan oil lens and Hamamatsu Photonics Orca-R2 camera. Unless otherwise stated in the figure legend, single planes of images are shown, but for quantification, 15 successive 400 nm slices were imaged to observe foci and NE-vacuole interactions near the edges of cells. Cells in Video 1 were imaged using a GE Healthcare Personal DeltaVision equipped with an Olympus IX71 microscope, 60 × NA 1.42 Plan Apochromat objective, and a Photometrics CoolSnap HQ2 camera. Time-lapse imaging was performed using an EMD Millipore CellASIC Onix Perfusion system with Y04C plates; successive Z stacks were imaged every 20 min for a total of 18 time points.

### Quantification of NE-vacuole interactions

FM4-64-stained cells that were imaged using 15 successive 400 nm Z stacks were examined to find the region of the cell, generally near the center, where the GFP-tagged nuclear rim was largest. Using this slice, the total length of the nuclear rim was calculated using the “freehand line” tool in Fiji. Then the length of the NE that directly contacted the NE was measured, again using the “freehand line” tool. The contact length was divided by the total length to quantify the amount of NE-vacuole interactions for a single cell.

### Electron microscopy

The glutaraldehyde/potassium permanganate protocol was performed largely as previously described (Wright, 2000). 20 ml of yeast were grown overnight to mid-log phase, then mixed with 20 ml of 2X fixation buffer (200 mM PIPES pH 6.8, 200 mM sorbitol, 2 mM MgCl_2_, 2 mM CaCl_2_, 4% glutaraldehye) with rocking for 5 min in a 50 ml conical tube. The cells were then centrifuged for 3 min at 3000 rpm and resuspended in 1X fixation buffer, where the cells were left overnight at 4°C while rocking. The following day, cells were washed 3 times for 10 min in 25 ml of ddH_2_0. Cells were transferred to 1.5 ml tubes and washed with 1 ml of freshly made 2% potassium permanganate for 5 min, then incubated with 1 ml 2% potassium permanganate for 1 hour while rocking. Cells were washed 5 times with 1 ml of ddH_2_0 for 1 min, transferred to new 1.5 ml tubes, then incubated with Uranyless (EMS, cat. no. 22409) for 1 hr while rocking. After washing cells 3 times with 1 ml of water for 1 min, the cells were dehydrated with increasing amounts of ethanol (25%, 50%, 75%, and 95%) for 5 min each. The cells were then washed 3 times with 100% ethanol for 5 min, then incubated overnight with 50% acetone/50% EMbed12 (EMS, cat. no. 14120) while rocking at RT. This was replaced with 100% EMmbed12 for 2 hours, which was repeated before finally resuspending cells in 500 μl EMmbed12 and the mixture was solidified by baking overnight at 60°C. Following sectioning, cells were stained with uranyl acetate and lead citrate before analyzing using a Philips/FEI T-12 transmission electron microscope.

### Cell lysates and western blotting

3-5 ml of cells were grown to mid-log phase at temperatures listed in figure legends, then centrifuged at 3,000 rpm in a tabletop centrifuge to pellet cells. Cells were transferred to 1.5 ml tubes and washed once with 1 ml YPD. After carefully removing all the remaining YPD, cell pellets were resuspended evenly in 40 μl 1X SDS sample buffer (60 mM Tris pH 6.8, 1% SDS, 10% glycerol, 1.67% β-mercaptoethanol, and 0.003% bromophenol blue) and immediately boiled for 5 min. Following boiling, acid-washed glass beads were added so that they encompassed the entire volume of liquid/cell suspension. Tubes were then vortexed heavily for 2 min. Finally, 60 μl 1X SDS sample buffer was added, the samples were quickly boiled for 30 sec, and then loaded onto polyacrylamide gels. SDS-PAGE and transfers were completed using established protocols and nitrocellulose membranes. Following transfer, membranes were stained with Ponceau S (0.1% Ponceau S, 5% acetic acid) to ensure protein levels were similar among samples. Ponceau S stain was removed by briefly washing membranes in 0.01N NaOH, which were then washed 3 times with 50 ml of water to remove residual NaOH. Membranes were blocked for 1 hour in TBST/7% milk, then incubated overnight at 4°C with primary antibodies used at 1:1,000 dilutions. The following day membranes were washed 3 times with 25 ml TBST for 5 min at RT, then incubated with TBST/7% milk for 1 hour at RT with 1:10,000 dilutions of secondary antibodies. Membranes were finally washed 3 times with 25 ml TBST for 5 min at RT, then imaged using a Li-Cor Odyssey system. Antibodies used include: affinity-purified polyclonal rabbit anti-Nup116 C-term (Iovine et al., 1995), monoclonal mouse anti-GFP (Millipore Sigma, cat. no. 11814460001 reconstituted at 0.4 mg/ml in water/0.05% sodium azide), monoclonal mouse anti-Pgk1 (ThermoFisher, cat. no. 459250), IRDye 680LT donkey anti-Mouse IgG (Li-Cor, cat. no. 926-68022), and IRDye 800CW Donkey anti-Rabbit IgG (Li-Cor, cat. no. 926-32213).

### NPC aggregation indices

Aggregation indices were quantified similarly to previous studies (Casey et al., 2015; Niepel et al., 2013). The average GFP-Nic96 fluorescence intensity at the nuclear rim was calculated by using the “freehand line” tool in Fiji to make a curved line that covered all of the rim in one plane of an image, which was then converted to quantifiable data using the “plot profile” tool; these numbers were averaged after background GFP intensity was subtracted. Average GFP-Nic96 fluorescence intensity was then subtracted from each plot profile measurement; the standard deviation of these absolute values was then divided by the average GFP-Nic96 fluorescence intensity to produce an aggregation index.

### Quantification of Nup85-GFP degradation

Li-Cor Image Studio was utilized to quantify the amounts of full-length Nup85-GFP (top band in anti-GFP blot in Fig. 5, G), as well as the three major GFP degradation products below full-length Nup85-GFP in *nup116ΔGLFG* strains. The amount of full-length Nup85-GFP was divided by the sum of the degradation products, then these ratios were compared to the value obtained from *nup116ΔGLFG* lysates. This was repeated 5 times from different independent experiments and was used to generate the data in Fig. 5, F.

### Serial dilutions

1 OD unit of yeast cells grown overnight at 25°C (unless temperature is otherwise noted in legends) to mid-log phase were pelleted via centrifugation and resuspended in 500 µl of sterile water, and were subsequently serially diluted five-fold 4 times in water. 4-5 µl was spotted onto plates, which were grown at temperatures listed in figure legends. Images were taken once wild type cells started to show growth at the greatest dilution.

### CPY secretion assay

Cells were serially diluted as described onto two YPD plates. One plate was covered with nitrocellulose to absorb secreted CPY, and both were grown overnight at 28°C. The nitrocellulose was removed and washed several times in water, while the plate without nitrocellulose was imaged. The nitrocellulose was then blocked for 1 hour in TBST/5% milk, and then incubated overnight at 4°C in 2 ml TBST/5% milk with a 1:1,000 dilution of anti-CPY Monoclonal antibody 10A5B5 (Invitrogen, A-6428). The nitrocellulose was then washed 3 times in 20 ml TBST at room temperature for 10 min, and incubated with 5 ml TBST/5% milk with a 1:10,000 dilution of IRDye 680LT donkey anti-Mouse IgG (Li-Cor, cat. no. 926-68022) for 1 hour at room temperature in a light-sensitive box. After washing 3 times with 20 ml TBST for 10 min at room temperature, the nitrocellulose was imaged on a Li-Cor Odyssey system.

### BODIPY 493/503 staining

1 ml of cells grown to mid-log phase in YPD were transferred to an amber 1.5 ml tube, along with 1 µl of 2 mM BODIPY 493/503/DMSO for a final concentration of 2 µM. Cells were incubated for 5 min at the temperature at which they were initially grown, washed two times with 1 ml YPD, then imaged.

### Quantification of NE-associated lipid droplets

The total amounts of LDs were measured FM4-64- and BODIPY-labeled in yeast strains by counting in max projections. In addition to staining LDs, BODIPY also weakly stains the NE and ER membranes, which is particularly evident when examining individual Z stacks. LDs at the NE were quantified by counting the amount at the NE that also co-labeled with FM4-64 in the same Z stack; this number was then divided by the total number of LDs.

### Quinacrine staining

1 ml of cells grown to mid-log phase in YPD were transferred to an amber 1.5 ml tube. The cells were centrifuged, and resuspended in YPD/50 mM sodium phosphate pH 7.6/100 μM quinacrine, then shaken at 36°C for 5 min. Cells were centrifuged and washed 2 times with pre-warmed YPD/50 mM sodium phosphate pH 7.6 before analyzing using fluorescence microscopy.

### Hoechst 33342 staining

Cells were incubated with 10 ug/ml Hoechst 33342 for 20 min in YPD amber tubes at specified temperatures, quickly washed twice with 1 ml YPD, then imaged.

### Calcofluor white staining

Cells were incubated for 5 min in 1 ml 0.5 µg/ml CCFW/YPD in amber tubes. Cells were then washed two times with 1 ml YPD, then imaged.

### Statistical Analysis

Data was depicted as mean with standard deviation error bars, except for bar and whiskers plots that were shown with mean and different percentiles. Unpaired two-tailed Student’s t-tests were used to compare GFP-Nic96, Nup85-GFP, and Nup85-GFP’ foci formation data, as well as western blot quantifications. For NE-vacuole interaction data, as well as data in Fig. 3 D, Fig. 6 C and Fig. S4 E, Dunn’s post-hoc test was utilized following a Kruskal–Wallis ANOVA. Mann-Whitney U tests were performed for data in Fig. 1 D and Fig. 3E. Microsoft Excel and Prism 8 were utilized to perform statistical tests. Data were considered significant if p-values were less than 0.05 for a particular statistical test.

## Supporting information

Strain list

## Acknowledgements

The authors wish to thank members of Susan Wente’s and Yi Ren’s laboratories for useful feedback about experiments and manuscript preparation, Patrick Lusk for discussions regarding NPC clustering experiments, Sepp D. Kohlwein for the *dga1Δ lro1Δ* strain, Kathleen Gould for use of her DeltaVision microscope and perfusion system, Alaina Willet for assistance with the DeltaVision, as well as Evan Krystofiak for assistance with EM sample preparation. EM experiments were performed in part using the Vanderbilt Cell Imaging Shared Resource (supported by NIH grants CA68485, DK20593, DK58404, DK59637 and EY08126). SRW and this work is supported by the NIGMS grant 5R37GM051219.

## Author Contributions

CLL and SRW cooperatively designed and interpreted experiments, and co-wrote the manuscript. CLL performed experiments.

## Competing interests

The authors declare no competing interests.

## Materials and Correspondence

Correspondence and reagent requests should be directed to Susan R. Wente using the email address.

## Abbreviations

NPC: nuclear pore complex Nup, nucleoporin
NE: nuclear envelope
NVJ: nucleus-vacuole junction
NTR: nuclear transport receptor
TEM: transmission electron microscopy LD, lipid droplet
SINC: storage of improperly assembled nuclear pore complexes compartment

---

**Video 1: GFP-Nic96 foci are not removed from the NE in *nup116ΔGLFG* cells**. *GFP-nic96 nup116ΔGLFG* cells grown overnight at 32°C, then imaged using a microfluidic chamber at 36°C for 4.5 hours. Frames are imaged 20 min apart, while max projections are shown.

**Supplementary Figure 1:**
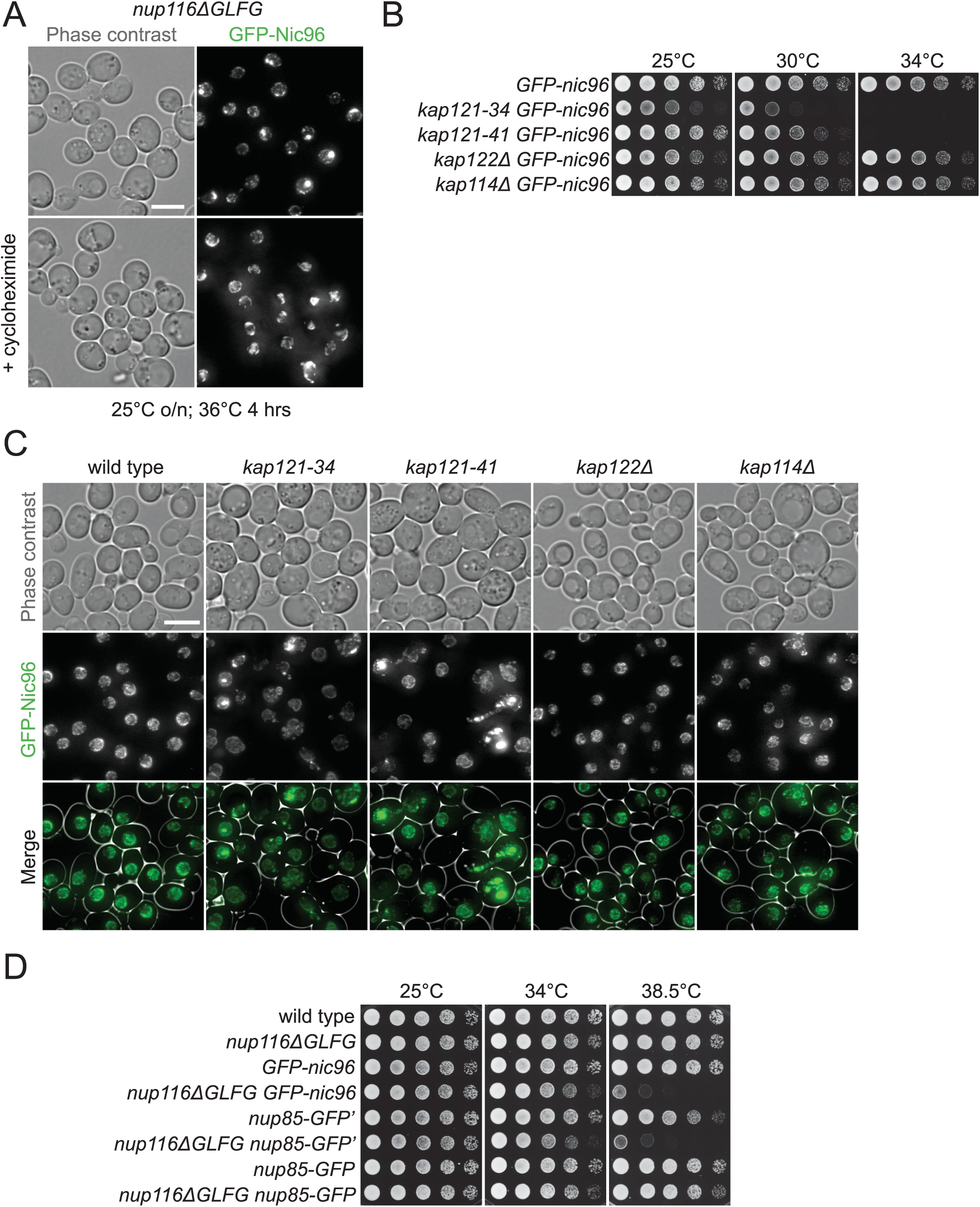
NPC assembly is disrupted in *nup116ΔGLFG* mutants, potentially due to inhibited Kap121-mediated transport. **(A)** Max projections of *GFP-nic96 nup116ΔGLFG* yeast that were grown to mid-log phase at 25°C, then shifted to 36°C for four hours in the absence (top) or presence (bottom) of 10 µg/ml cycloheximide. Bar, 5 µm. **(B)** Serial dilutions of different *GFP-nic96 kap* mutants on YPD plates. **(C)** Max projections of *GFP-nic96* yeast strains grown to at 32°C. Bar, 5 µm. **(D)** Serial dilutions of the listed strains on YPD plates.

**Supplementary Figure 2:**
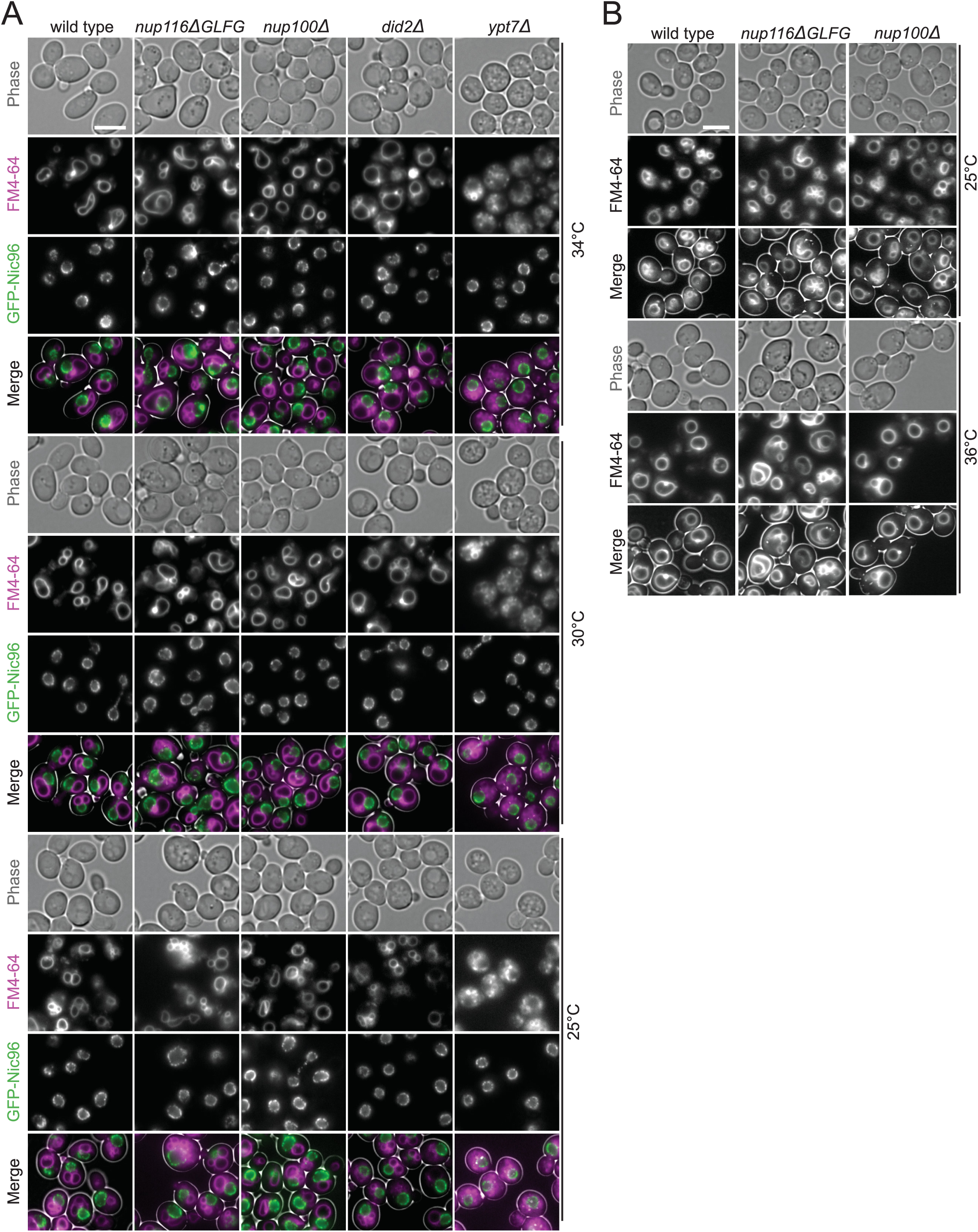
Images of *GFP-nic96* strains at 25, 30, and 34°C. **(A)** FM4-64-stained *GFP-nic96* strains grown at 25, 30, or 34°C that were used to quantify data in Fig. 1, B-D. Bar, 5 µm. **(B)** FM4-64-stained wild type, *nup116ΔGLFG*, and *nup100Δ* cells grown at 25 or 36°C; the similar vacuole morphology phenotypes indicate NE-vacuole contacts are not significantly impacted by *GFP-nic96*.

**Supplementary Figure 3:**
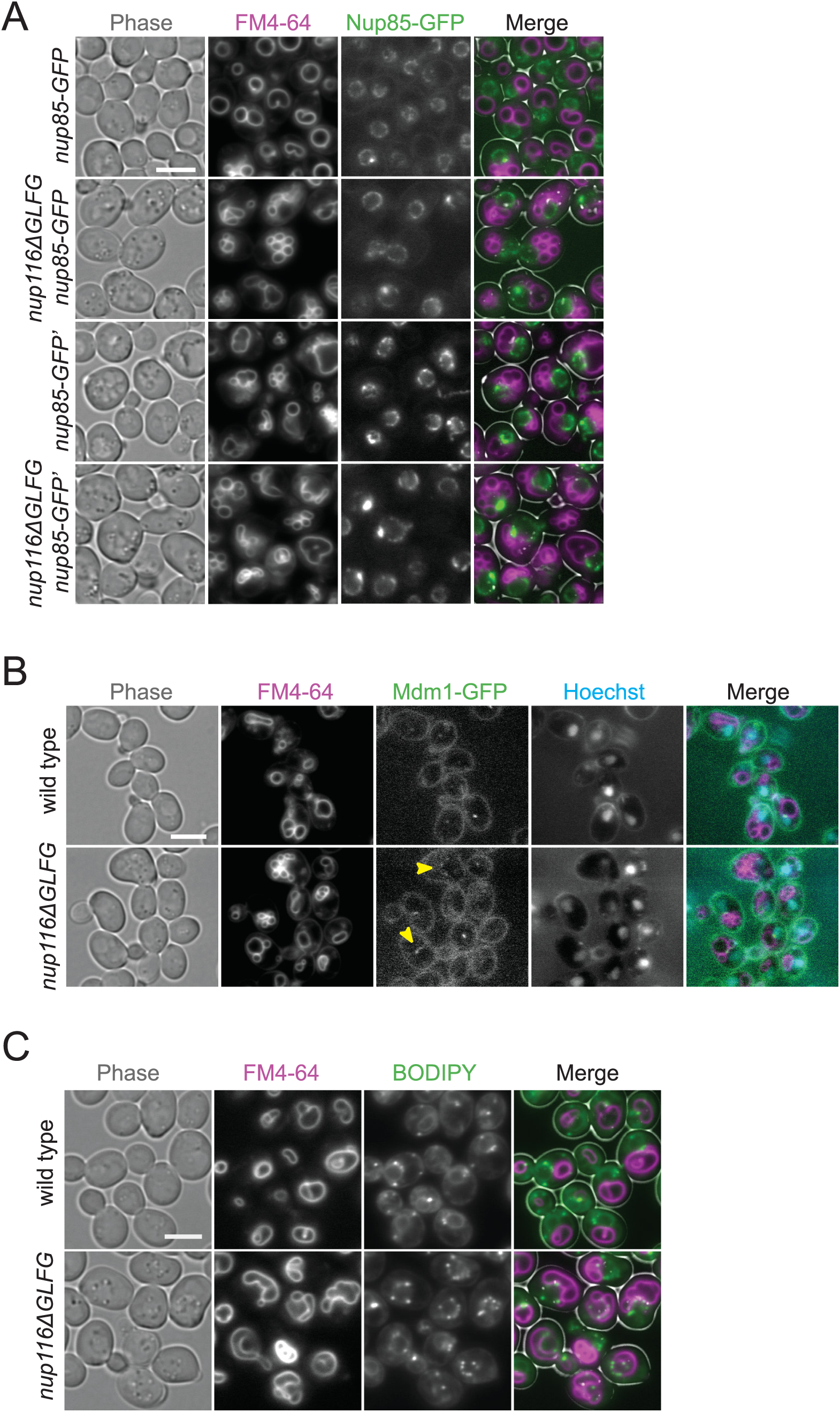
Lipid droplets and Mdm1 are enriched at NE-vacuole interfaces in *nup116ΔGLFG* cells. **(A)** FM4-64-stained *nup85-GFP* or *nup85-GFP’* strains grown at 30°C; these images were utilized to quantify data in Fig. 2 F. Bar, 5 µm. **(B)** FM4-64- and Hoechst-stained *mdm1-GFP* strains grown at 30°C. Arrows point to cells where Mdm1-GFP localizes around multiple regions of NE-vacuole contacts. Bar, 5 µm. **(C)** Images of FM4-64- and BODIPY-stained wild type and *nup116ΔGLFG* cells grown at 34°C. Bar, 5 µm. These cells, as well as other independent experiments, were used to quantify data in Fig. 3 E.

**Supplementary Figure 4:**
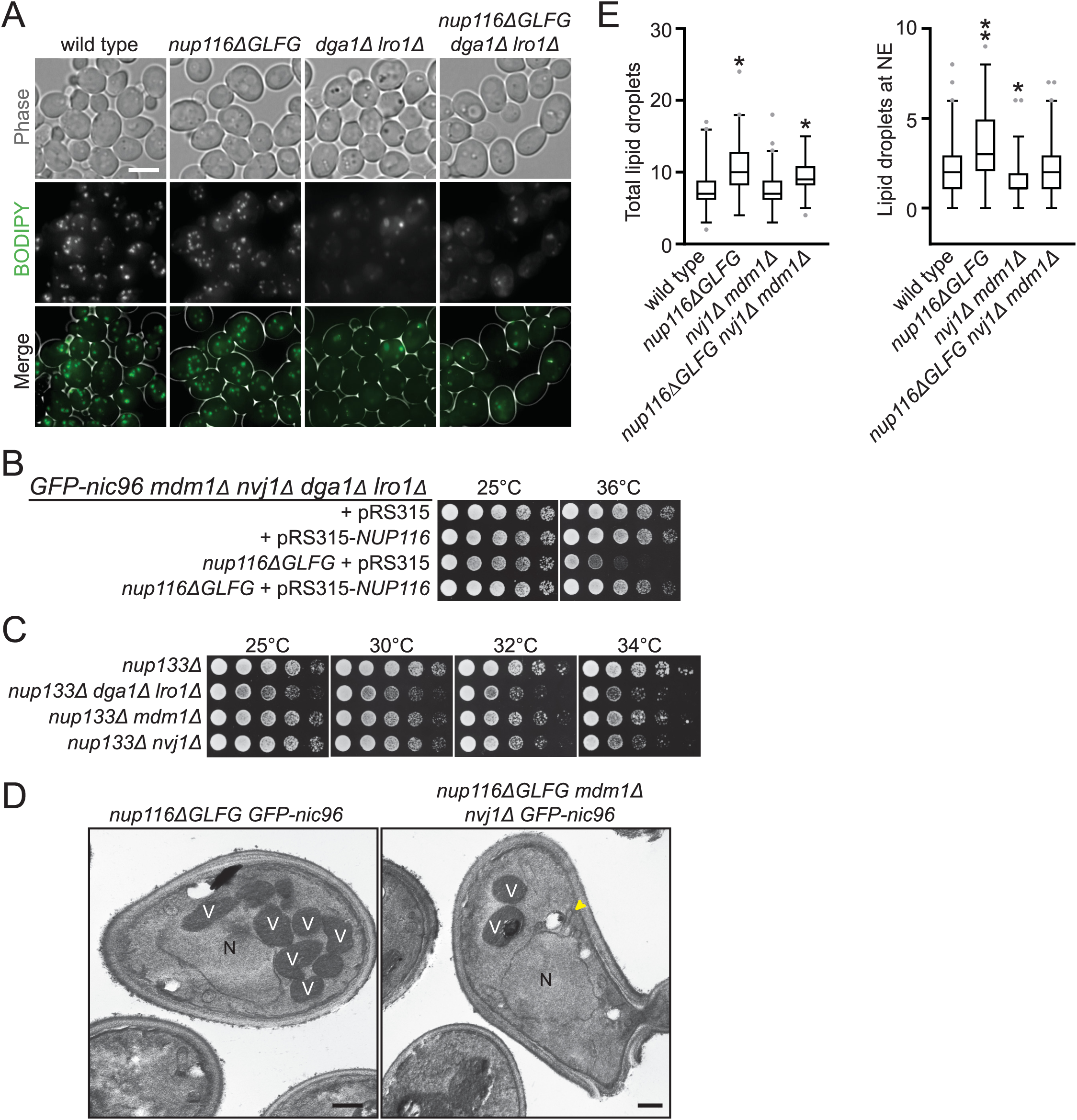
NVJs impact the viability of *nup133Δ* cells, as well as the localization of LDs and NE morphology in *nup116ΔGLFG* strains. **(A)** BODIPY-stained strains grown at 30°C. Bar, 5 µm. **(B)** Serial dilutions of *GFP-nic96 mdm1Δ nvj1Δ dga1Δ lro1Δ* or *nup116ΔGLFG GFP-nic96 mdm1Δ nvj1Δ dga1Δ lro1Δ* cells transformed with pRS315-*NUP116* or pRS315, grown in SC-leu and then plated onto YPD plates. **(C)** Serial dilutions of the listed strains on YPD plates. **(D)** TEM images of cells grown at 30°C. Nuclei are labeled with a black “N”, while heavily stained vacuoles are labeled with a white “V”. Arrow points to NE herniation. Bars, 500 nm. **(E)** Quantification of total LD number (left) and LDs at the NE (right) in the listed strains grown at 34°C. Box and whiskers represent 2.5-97.5 percentile. Asterisk indicates a p-value of less than 0.01 compared to wild type using Dunn’s post-hoc test, while two asterisks indicate a p-value of less than 0.02 when also compared to *nup116ΔGLFG mdm1Δ nvj1Δ*.

**Supplementary Figure 5:**
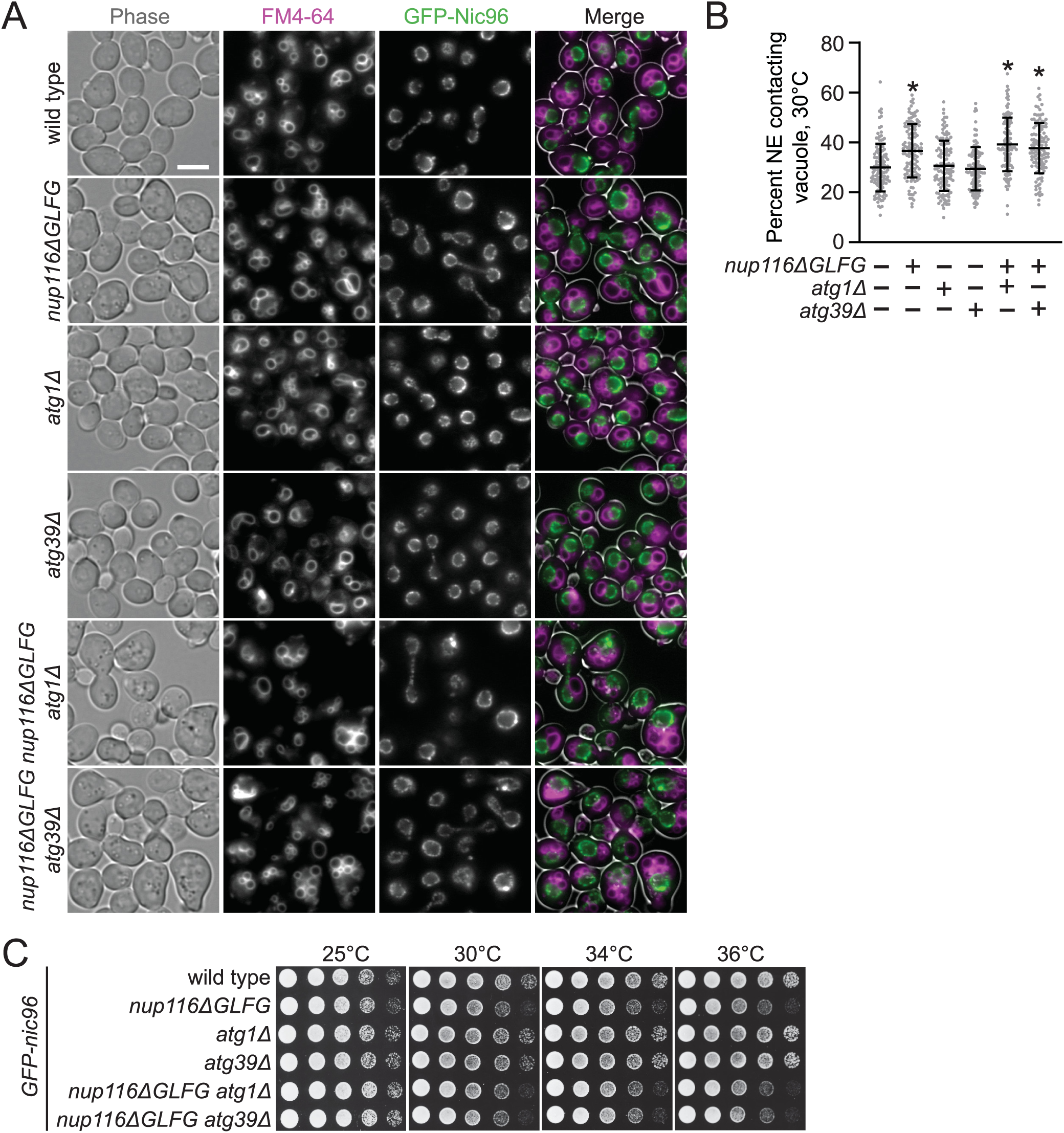
*ATG1* and ATG39 do not impact NE-vacuole interactions in *nup116ΔGLFG* mutants or their viability. **(A)** FM4-64-stained *GFP-nic96* strains grown at 30°C. Bar, 5 µm. **(B)** Quantification of NE-vacuole interactions in the listed *GFP-nic96* strains at 30°C. Each dot represents value from one cell, and 40 cells from 3 independent experiments were analyzed. Asterisk indicates a p-value of less than 0.01 when compared to wild type using Dunn’s post-hoc test. Error bars show standard deviation. **(C)** Serial dilutions of the listed *GFP-nic96* strains on YPD plates.

