## Supplementary material for "Nuclear envelope-vacuole contacts mitigate nuclear pore complex assembly stress": Strain list

| Strain number | Genotype | Source |
|---|---|---|
| BY4741 | *MATa his3Δ1 leu2Δ0 met15Δ0 ura3Δ0* | Brachmann, C. B. et al. |
| BY4742 | *MATα his3Δ1 leu2Δ0 lys2Δ0 ura3Δ0* | Brachmann, C. B. et al. |
| *nup85-GFP* | *MATa his3Δ1 leu2Δ0 met15Δ0 ura3Δ0 nup85-GFP::HIS3 (nup85-GFP')* | Huh, W.-K. et al. |
| SWY2791 | *MATα trp1-1 ura3-1 leu2-3,112 his3-11,15 nup116ΔGLFG* | Strawn, L. et al. |
| SWY6248 | *MATa his3Δ1 leu2Δ0 met15Δ0 ura3Δ0 nup100Δ::KAN* | Lord, C. L., Ospovat, O. & Wente, S. R. |
| SWY6615 | *MATα his3Δ1 leu2Δ0 lys2Δ0 ura3Δ0 nup116ΔGLFG arg4Δ::KAN* | This manuscript |
| SWY6743 | *MATα his3Δ1 leu2Δ0 lys2Δ0 ura3Δ0 ypt7Δ::KAN* | This manuscript |
| SWY6746 | *MATα his3Δ1 leu2Δ0 lys2Δ0 ura3Δ0 did2Δ::KAN* | This manuscript |
| SWY6756 | *MATa his3Δ1 leu2Δ0 met15Δ0 ura3Δ0 GFP-nic96::HIS3* | This manuscript |
| SWY6757 | *MATα his3Δ1 leu2Δ0 lys2Δ0 ura3Δ0 nup116ΔGLFG arg4Δ::KAN GFP-nic96::HIS3* | This manuscript |
| SWY6758 | *MATa his3Δ1 leu2Δ0 met15Δ0 ura3Δ0 nup100Δ::KAN GFP-nic96::HIS3* | This manuscript |
| SWY6759 | *MATα his3Δ1 leu2Δ0 lys2Δ0 ura3Δ0 did2Δ::KAN GFP-nic96::HIS3* | This manuscript |
| SWY6760 | *MATα his3Δ1 leu2Δ0 lys2Δ0 ura3Δ0 ypt7Δ::KAN GFP-nic96::HIS3* | This manuscript |
| SWY6786 | *MATα his3Δ1 leu2Δ0 lys2Δ0 ura3Δ0 nup116ΔGLFG arg4Δ::KAN nup85-GFP::HYG* | This manuscript |
| SWY6813 | *MATa his3Δ1 leu2Δ0 met15Δ0 ura3Δ0 nup120Δ::KAN* | This manuscript |
| SWY6814 | *MATa his3Δ1 leu2Δ0 met15Δ0 ura3Δ0 nup133Δ::KAN* | This manuscript |
| SWY6815 | *MATa his3Δ1 leu2Δ0 met15Δ0 ura3Δ0 vps4 Δ ::KAN GFP-nic96::HIS3* | This manuscript |
| SWY6816 | *MATa his3Δ1 leu2Δ0 met15Δ0 ura3Δ0 nup120Δ::KAN GFP-nic96::HIS3* | This manuscript |
| SWY6817 | *MATa his3Δ1 leu2Δ0 met15Δ0 ura3Δ0 nup133Δ::KAN GFP-nic96::HIS3* | This manuscript |
| SWY6824 | *MATa his3Δ1 leu2Δ0 met15Δ0 ura3Δ0 kap121-34::KAN GFP-nic96::HIS3* | This manuscript |
| SWY6825 | *MATa his3Δ1 leu2Δ0 met15Δ0 ura3Δ0 kap121-41::KAN GFP-nic96::HIS3* | This manuscript |
| SWY6826 | *MATa his3Δ1 leu2Δ0 met15Δ0 ura3Δ0 kap122Δ::KAN GFP-nic96::HIS3* | This manuscript |
| SWY6827 | *MATa his3Δ1 leu2Δ0 met15Δ0 ura3Δ0 kap114Δ::KAN GFP-nic96::HIS3* | This manuscript |
| SWY6828 | *MATa his3Δ1 leu2Δ0 met15Δ0 ura3Δ0 chm7 Δ ::KAN GFP-nic96::HIS3* | This manuscript |
| SWY6832 | *his3Δ1 leu2Δ0 ura3Δ0 vps4 Δ ::KAN nup116 Δ GLFG GFP-nic96::HIS3* | This manuscript |
| SWY6867 | *his3Δ1 leu2Δ0 ura3Δ0 nup116ΔGLFG nup85-GFP::HIS (nup85-GFP')* | This manuscript |
| SWY6868 | *his3Δ1 leu2Δ0 ura3Δ0 nup85-GFP::HYG* | This manuscript |
| SWY6872 | *his3Δ1 leu2Δ0 ura3Δ0 chm7 Δ ::KAN nup116 Δ GLFG GFP-nic96::HIS3* | This manuscript |
| SWY6877 | *his3Δ1 leu2Δ0 ura3Δ0 pho88-GFP::HIS3 nup120Δ::KAN* | This manuscript |
| SWY6878 | *his3Δ1 leu2Δ0 ura3Δ0 pho88-GFP::HIS3 nup133Δ::KAN* | This manuscript |
| SWY6885 | *his3Δ1 leu2Δ0 ura3Δ0 nup116ΔGLFG nup120Δ::KAN* | This manuscript |
| SWY6885 | *his3Δ1 leu2Δ0 ura3Δ0 vps4 Δ ::KAN vps4 Δ::KAN GFP-nic96::HIS3* | This manuscript |
| SWY6887 | *his3Δ1 leu2Δ0 ura3Δ0 nup116ΔGLFG nup133Δ::KAN* | This manuscript |
| SWY6889 | *his3Δ1 leu2Δ0 ura3Δ0 nvj1Δ::HYG mdm1Δ::HYG GFP-nic96::HIS3* | This manuscript |
| SWY6890 | *his3Δ1 leu2Δ0 ura3Δ0 nvj1Δ::HYG mdm1Δ::HYG nup116ΔGLFG GFP-nic96::HIS3* | This manuscript |
| SWY6894 | *his3Δ1 leu2Δ0 ura3Δ0 pep4 Δ ::KAN pho88-GFP::HIS3* | This manuscript |
| SWY6895 | *his3Δ1 leu2Δ0 ura3Δ0 pep4 Δ ::KAN pho88-GFP::HIS3 nup116 Δ GLFG* | This manuscript |
| SWY6896 | *his3Δ1 leu2Δ0 ura3Δ0 pep4 Δ ::KAN nup85-GFP::HYG* | This manuscript |
| SWY6897 | *his3Δ1 leu2Δ0 ura3Δ0 pep4 Δ ::KAN nup85-GFP::HYG nup116 Δ GLFG* | This manuscript |
| SWY6898 | *his3Δ1 leu2Δ0 ura3Δ0 dga1Δ::KAN lro1Δ::KAN nup133Δ::KAN* | This manuscript |
| SWY6899 | *his3Δ1 leu2Δ0 ura3Δ0 dga1Δ::KAN lro1Δ::KAN nup116ΔGLFG::KAN* | This manuscript |
| SWY6900 | *MATα his3Δ1 leu2Δ0 lys2Δ0 ura3Δ0 dga1Δ::KAN lro1Δ::KAN GFP-nic96::HIS3* | This manuscript |
| SWY6901 | *his3Δ1 leu2Δ0 ura3Δ0 dga1Δ::KAN lro1Δ::KAN nup116ΔGLFG::KAN GFP-nic96::HIS3* | This manuscript |
| SWY6902 | *MATa his3Δ1 leu2Δ0 met15Δ0 ura3Δ0 nvj1-GFP::HIS3 (check if hyg or his)* | This manuscript |
| SWY6903 | *his3Δ1 leu2Δ0 ura3Δ0 nup116ΔGLFG nvj1-GFP::HIS3 (check if hyg or his)* | This manuscript |
| SWY6906 | *MATa his3Δ1 leu2Δ0 met15Δ0 ura3Δ0 nup133Δ::KAN mdm1Δ::HYG* | This manuscript |
| SWY6911 | *MATa his3Δ1 leu2Δ0 met15Δ0 ura3Δ0 mdm1-GFP::HYG* | This manuscript |
| SWY6912 | *his3Δ1 leu2Δ0 ura3Δ0 nup116ΔGLFG mdm1-GFP::HYG* | This manuscript |
| SWY6913 | *his3Δ1 leu2Δ0 ura3Δ0 dga1Δ::KAN lro1Δ::KAN nup85-GFP::HIS3* | This manuscript |
| SWY6914 | *his3Δ1 leu2Δ0 ura3Δ0 dga1Δ::KAN lro1Δ::KAN nup85-GFP::HIS3 nup116 Δ GLFG* | This manuscript |
| SWY6915 | *his3Δ1 leu2Δ0 ura3Δ0 nvj1Δ::HYG mdm1Δ::HYG nup85-GFP::HIS3* | This manuscript |
| SWY6916 | *his3Δ1 leu2Δ0 ura3Δ0 nvj1Δ::HYG mdm1Δ::HYG nup85-GFP::HIS3 nup116 Δ GLFG* | This manuscript |
| SWY6917 | *his3Δ1 leu2Δ0 ura3Δ0 nup133Δ::KAN nvj1Δ::HYG* | This manuscript |
| SWY6923 | *his3Δ1 leu2Δ0 ura3Δ0 dga1Δ::KAN lro1Δ::KAN mdm1::HYG nvj1::HYG GFP-nic96::HIS3* | This manuscript |
| SWY6924 | *his3Δ1 leu2Δ0 ura3Δ0 dga1Δ::KAN lro1Δ::KAN mdm1::HYG nvj1::HYG nup116ΔGLFG GFP-nic96::HIS3* | This manuscript |
| SWY6939 | *MATa his3Δ1 leu2Δ0 met15Δ0 ura3Δ0 GFP-nic96::HIS3 atg1 Δ ::HYG* | This manuscript |
| SWY6940 | *MATa his3Δ1 leu2Δ0 met15Δ0 ura3Δ0 GFP-nic96::HIS3 atg39 Δ ::HYG* | This manuscript |
| SWY6941 | *MATα his3Δ1 leu2Δ0 lys2Δ0 ura3Δ0 nup116ΔGLFG arg4Δ::KAN GFP-nic96::HIS3 atg1 Δ ::HYG* | This manuscript |
| SWY6942 | *MATα his3Δ1 leu2Δ0 lys2Δ0 ura3Δ0 nup116ΔGLFG arg4Δ::KAN GFP-nic96::HIS3 atg39 Δ ::HYG* | This manuscript |
| SWY6947 | *his3Δ1 leu2Δ0 ura3Δ0 atg1 Δ ::KAN nup85-GFP::HYG* | This manuscript |
| SWY6948 | *his3Δ1 leu2Δ0 ura3Δ0 atg1 Δ ::KAN nup85-GFP::HYG nup116 Δ GLFG* | This manuscript |
| SWY6949 | *his3Δ1 leu2Δ0 ura3Δ0 atg1 Δ ::HIS3 nup85-GFP::HYG nup116 Δ GLFG mdm1::HYG nvj1::HYG* | This manuscript |
| *vph1 Δ* | *MATa his3Δ1 leu2Δ0 met15Δ0 ura3Δ0 vph1Δ::KAN* | Giaever, G. et al. |
| YJP1075 | *MATα his3Δ1 leu2Δ0 lys2Δ0 ura3Δ0 dga1Δ::KAN lro1Δ::KAN* | Kohlwein, S. D. et al. |
